# Activation of intestinal immunity by pathogen effector-triggered aggregation of lysosomal TIR-1/SARM1

**DOI:** 10.1101/2023.12.04.569946

**Authors:** Samantha Y. Tse-Kang, Khursheed A. Wani, Nicholas D. Peterson, Amanda Page, Read Pukkila-Worley

**Affiliations:** Program in Innate Immunity, Division of Infectious Diseases and Immunology, Department of Medicine, University of Massachusetts Chan Medical School, Worcester, MA 01655, United States of America

**Keywords:** effector-triggered <u>immunity</u>, lysosome-related organelles, intestinal immunity, TIR-1/SARM1, *Pseudomonas aeruginosa*, *Caenorhabditis elegans*, phenazines

## Abstract

TIR-domain proteins with enzymatic activity are essential for immunity in plants, animals, and bacteria. However, it is not known how these proteins function in pathogen sensing in animals. We discovered that a TIR-domain protein (TIR-1/SARM1) is strategically expressed on the membranes of a lysosomal sub-compartment, which enables intestinal epithelial cells in the nematode *C. elegans* to survey for pathogen effector-triggered host damage. We showed that a redox active virulence effector secreted by the bacterial pathogen *Pseudomonas aeruginosa* alkalinized and condensed a specific subset of lysosomes by inducing intracellular oxidative stress. Concentration of TIR-1/SARM1 on the surface of these organelles triggered its multimerization, which engages its intrinsic NADase activity, to activate the p38 innate immune pathway and protect the host against microbial intoxication. Thus, lysosomal TIR-1/SARM1 is a sensor for oxidative stress induced by pathogenic bacteria to activate metazoan intestinal immunity.

## INTRODUCTION

Toll/interleukin-1/resistance gene (TIR)-domain proteins are conserved across the tree of life and are essential for innate immunity. Toll-like receptors (TLRs) and interleukin-1 receptors (IL-1Rs) in animals, and nucleotide-binding leucine-rich repeat receptors (NLRs) in plants all contain TIR domains necessary for their function in pathogen recognition and immune activation.^1–3^ In plants, specialized TIR domain-containing proteins are enzymes that metabolize nicotinamide adenine dinucleotide (NAD+) to synthesize immuno-regulatory, second messenger metabolites that coordinate anti-pathogen defenses.^4^ Likewise, bacterial TIR is an NAD^+^ hydrolase enzyme required for defense against phage infection.^5^ Despite their evolutionarily ancient functions in innate immunity, it is not known how TIR enzymes function in animal innate immunity.

The soil-dwelling nematode *Caenorhabditis elegans* encodes two proteins with a TIR domain, one of which (TIR-1) is an NAD^+^ hydrolase (NADase) enzyme. *C. elegans* TIR-1, the nematode homolog of mammalian sterile alpha and TIR motif-containing 1 (SARM1), is essential for innate immunity, functioning as the upstream activator of the conserved p38 PMK-1 mitogen-activated protein kinase (MAPK) host defense pathway.^6–11^ We previously showed TIR-1/SARM1 multimerizes into protein assemblies (puncta) in intestinal epithelial cells.^12^ Functionally, concentration of TIR-1/SARM1 in this manner engages its intrinsic NADase activity, which activates p38 PMK-1 signaling to promote resistance to pathogen infection.^12^ Intriguingly, multimerization of plant TIR is also a prerequisite for its NADase activity,^13,14^ as it is for human SARM1, which promotes axonal degeneration following neuronal injury.^15,16^ In this study, we sought to determine how pathogen infection initiates TIR-1/SARM1 multimerization and its intrinsic NADase activity to promote host defense against bacterial infection in the animal intestine.

Nematodes live in microbe-rich habitats and consume bacteria for nutrition. *C. elegans* are therefore constantly exposed to structural features of bacteria that are sensed by pattern recognition receptors. As such, nematodes lost canonical mechanisms of bacterial pattern recognition during evolution.^17–19^ Instead, *C. elegans* evolved strategies to survey for so-called patterns of pathogenesis, rather than structural features of bacteria, to distinguish infectious pathogens from its bacterial food. For example, *C. elegans* uses a non-canonical pattern recognition system to intercept pathogen-derived metabolite signatures of bacterial growth and virulence to evaluate the threat posed by environmental pathogens.^20^ Additionally, pathogen-induced disruption of host translation,^21–23^ mitochondrial respiration,^24–26^ and the proteasome,^27,28^ as well as cell damage^29^ and physical distention of the intestinal lumen,^30–32^ activate innate immune defenses. In this context, the conservation of a TIR protein with NADase activity in innate immunity is noteworthy in an animal that does not utilize TIR domain-containing TLRs as pattern recognition receptors. This observation suggests that TIR enzymes are fundamental for the evolution of innate immunity in general and may be involved in sensing bacterial pattern of pathogenesis to activate innate immunity.

Here, we discovered that TIR-1/SARM1 puncta localized to the membranes of a specific lysosomal sub-compartment called lysosome-related organelles. Oxidative stress induced by pyocyanin, a redox active virulence effector secreted by the bacterial pathogen *Pseudomonas aeruginosa*, alkalinized and condensed lysosome-related organelles in the intestinal epithelial cells of *C. elegans*. The changes in this lysosomal sub-compartment induced the multimerization of TIR-1/SARM1 on the membrane of these organelles, which engages the intrinsic NADase activity of the protein complex, to initiate p38 PMK-1 immune pathway signaling. These data characterize a mechanism of enzymatic TIR activation of animal immunity and demonstrate that a specific lysosomal compartment is a dynamic scaffold for TIR-1/SARM1 that enables intestinal epithelial cells to survey for the effects of a bacterial pathogen to activate innate immune defenses.

## RESULTS

### TIR-1/SARM1 is expressed on lysosome-related organelles in *C. elegans* intestinal epithelial cells

Previously, we used clustered regularly interspaced short palindromic repeats (CRISPR)-Cas9 to engineer a *C. elegans* strain that expresses TIR-1/SARM1 protein tagged with the fluorophore wrmScarlet at its endogenous genomic locus.^12^ We observed that pathogen infection induces TIR-1::wrmScarlet multimerization into large protein assemblies, or puncta, in the intestinal epithelium.^12^ Here, we sought to characterize the mechanism by which infection promoted TIR-1/SARM1 assembly to activate the p38 PMK-1 innate immune pathway.

The dim fluorescence of endogenous TIR-1::wrmScarlet expression and autofluorescence in wild-type *C. elegans* intestinal epithelial cells prevented detailed characterization of the sub-cellular localization of TIR-1/SARM1 in live animals. We therefore performed immunostaining on fixed animals to visualize TIR-1/SARM1 in intestinal tissues. For these studies, a *C. elegans* strain that we previously generated using CRISPR-Cas9, which contains TIR-1/SARM1 tagged at its endogenous locus with a 3xFLAG sequence,^12^ was probed with an anti-FLAG antibody in fixed samples. Immunostaining of TIR-1/SARM1 confirmed that this protein formed puncta in intestinal epithelial cells (**Figure 1A**). Interestingly, we also observed that a subset of TIR-1/SARM1 puncta formed vesicular structures in the intestine (**Figure 1A**, dashed arrows), which were not visualized either in animals treated with RNA interference (RNAi) specific for *tir-1* (**Figure 1B**) or in untagged animals (**Figure S1A**).

**Figure 1.**
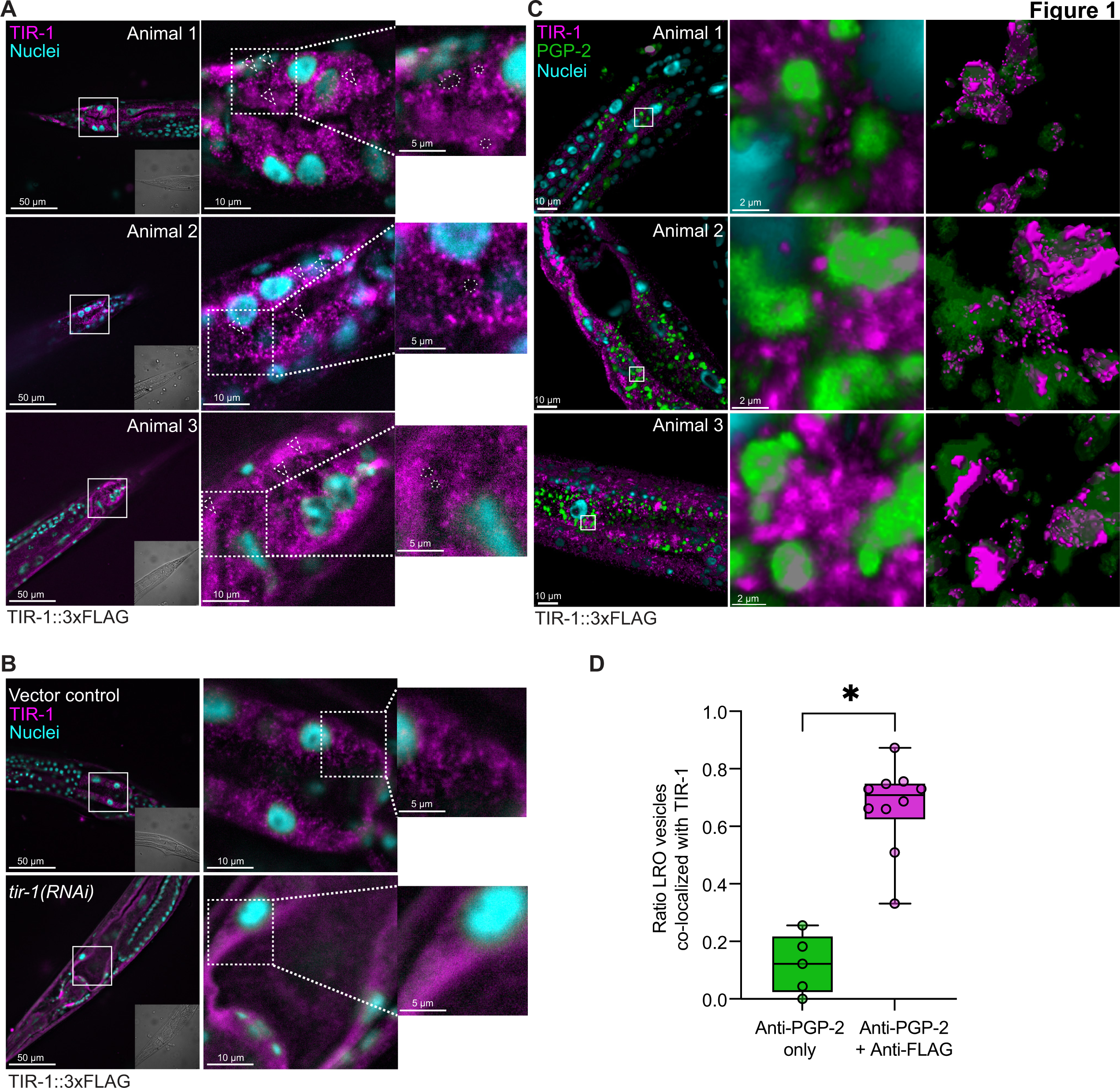
TIR-1/SARM1 is expressed on lysosome-related organelles in *C. elegans* intestinal epithelial cells. **(A)** Representative images of three fixed *C. elegans* TIR-1::3xFLAG animals at the L4 larval stage immunostained with an anti-FLAG antibody (for TIR-1) and DAPI. Insets in the left panel represent corresponding DIC images. Dotted boxes indicate higher magnifications. Dashed arrows in the middle panel indicate multiple vesicular structures. Dotted circles in the right panel highlight lumen of vesicular structures. **(B)** Vector- and *tir-1(RNAi)*-treated *C. elegans* TIR-1::3xFLAG animals were immunostained with an anti-FLAG antibody (for TIR-1) and DAPI. **(C)** Representative images of three *C. elegans* TIR-1::3xFLAG animals co-immunostained with antibodies against FLAG (for TIR-1) or PGP-2 (for lysosome-related organelles). Images are three-dimensional reconstructions of Z-stacks. The last column shows TIR-1/SARM1 (magenta) co-localized with PGP-2+ lysosome-related organelles (green). **(D)** Quantification of TIR-1/SARM1 co-localization on PGP-2+ lysosome-related organelles (LRO) in animals stained only with anti-PGP-2 (control) or co-immunostained with both anti-FLAG and anti-PGP-2 by Aivia (Leica). Scale bars as indicated. Source data for this figure is in Table S2. See also Fig. S1. *equals p<0.05 (unpaired t-test).

In a companion manuscript that we co-submitted with this paper, we discovered that p38 PMK-1 propagated its own activation by promoting aggregation and activation of its upstream regulator, TIR-1/SARM1. We performed a forward genetic screen to determine how the p38 positive feedback loop is regulated to prevent the deleterious effects of unchecked immune activation and identified a previously uncharacterized gene Y42G9A.1, which we named regulator *o*f *<u>tir</u>-1* (*rotr-1*). We found that ROTR-1 functioned in the intestine to suppress TIR-1/SARM1 multimerization and to maintain the size of lysosome-related organelles. Therefore, we hypothesized that TIR-1/SARM1 is expressed on the surface of lysosome-related organelles to control p38 PMK-1 pathway activation.

By immunostaining, we found that the vesicles that expressed TIR-1/SARM1 were lysosome-related organelles. Specifically, we co-immunostained *C. elegans* TIR-1::3xFLAG animals with an anti-FLAG antibody and a previously generated anti-PGP-2 antibody (**Figure 1C and Supplemental Video 1**).^33^ PGP-2 is an ATP-binding cassette transporter that is specifically expressed on the membranes of lysosome-related organelles.^33^ Three-dimensional reconstruction of z-stacked images from co-immunostained animals revealed that TIR-1/SARM1 localized to the surface of lysosome-related organelles (**Figure 1C and Supplemental Video 1**). We used the machine-learning software Aivia (Leica) to quantify the co-localization between TIR-1/SARM1+ and PGP-2+ vesicles in three dimensions (see Materials and Methods). Of all PGP-2+ vesicles (green in **Figure 1C**), TIR-1/SARM1 (magenta in **Figure 1C**) co-localized with a median of 70.8% of the vesicles (**Figure 1D**). As a control, we used Aivia to quantify the co-localization of PGP-2+ vesicles and non-specific Alexa Fluor 555+ signal in animals that were incubated with both anti-mouse and anti-rabbit secondary antibodies, but stained only with the anti-PGP-2 antibody. We observed a significant difference in the co-localization of green and magenta signals between this control and the treatment conditions, as quantified by Aivia (**Figure 1D**). Finally, we confirmed the validity of the anti-PGP-2 antibody in detecting lysosome-related organelles within intestinal epithelial cells by performing immunostaining on a *C. elegans* strain that carries a translational reporter for PGP-2 (PGP-2::GFP), which demonstrated that PGP-2::GFP fluorescence co-localized with staining by the anti-PGP-2 antibody (**Figure S1B**). We also confirmed that the *pgp-2* loss-of-function mutants had abrogated staining by the anti-PGP-2 antibody (**Figure S1C**).

In summary, our data demonstrate that TIR-1/SARM1 is expressed on lysosome-related organelles in the *C. elegans* intestine.

### TIR-1/SARM1 co-localization with lysosome-related organelles can be visualized in live *C. elegans*

As an orthologous means to confirm that TIR-1/SARM1 is expressed on lysosome-related organelles, we imaged TIR-1::wrmScarlet under an activating condition, which increased its overall expression and enabled co-localization experiments with GFP-labeled cellular compartments. In a companion manuscript, we found that the p38 PMK-1 pathway potentiated its own activation in a positive feedforward loop by increasing the expression and aggregation of TIR-1/SARM1. RNAi-mediated knockdown of *vhp-1,* the phosphatase that dephosphorylates p38 PMK-1, caused robust induction of both TIR-1::3xFLAG and TIR-1::wrmScarlet expression (see companion manuscript).

We utilized the increased expression of TIR-1::wrmScarlet conferred by *vhp-1(RNAi)* treatment to perform co-localization experiments. We crossed *C. elegans* TIR-1::wrmScarlet with strains that express GFP translational fusion proteins – PGP-2 (PGP-2::GFP), which marks lysosome-related organelles, and LMP-1 (LMP-1::GFP), which marks lysosomes.^34,35^ Treatment of wild-type animals with *vhp-1(RNAi)* revealed that TIR-1::wrmScarlet formed puncta on the membranes of vesicles that expressed PGP-2::GFP (**Figure 2A**, see filled arrows in higher magnification image), but not LMP-1::GFP (**Figure S2**). Analysis using ImageJ and the JACoP plug-in demonstrated that TIR-1::wrmScarlet significantly co-localized with PGP-2::GFP (*r* = 0.426, p<0.05, Pearson correlation coefficient) (**Figure 2B**), but not LMP-1::GFP (p=not significant, Pearson correlation coefficient) (**Figure 2C**). RNAi-mediated knockdown of *tir-1* suppressed TIR-1::wrmScarlet puncta formation (**Figures 2A and 2B**).

**Figure 2.**
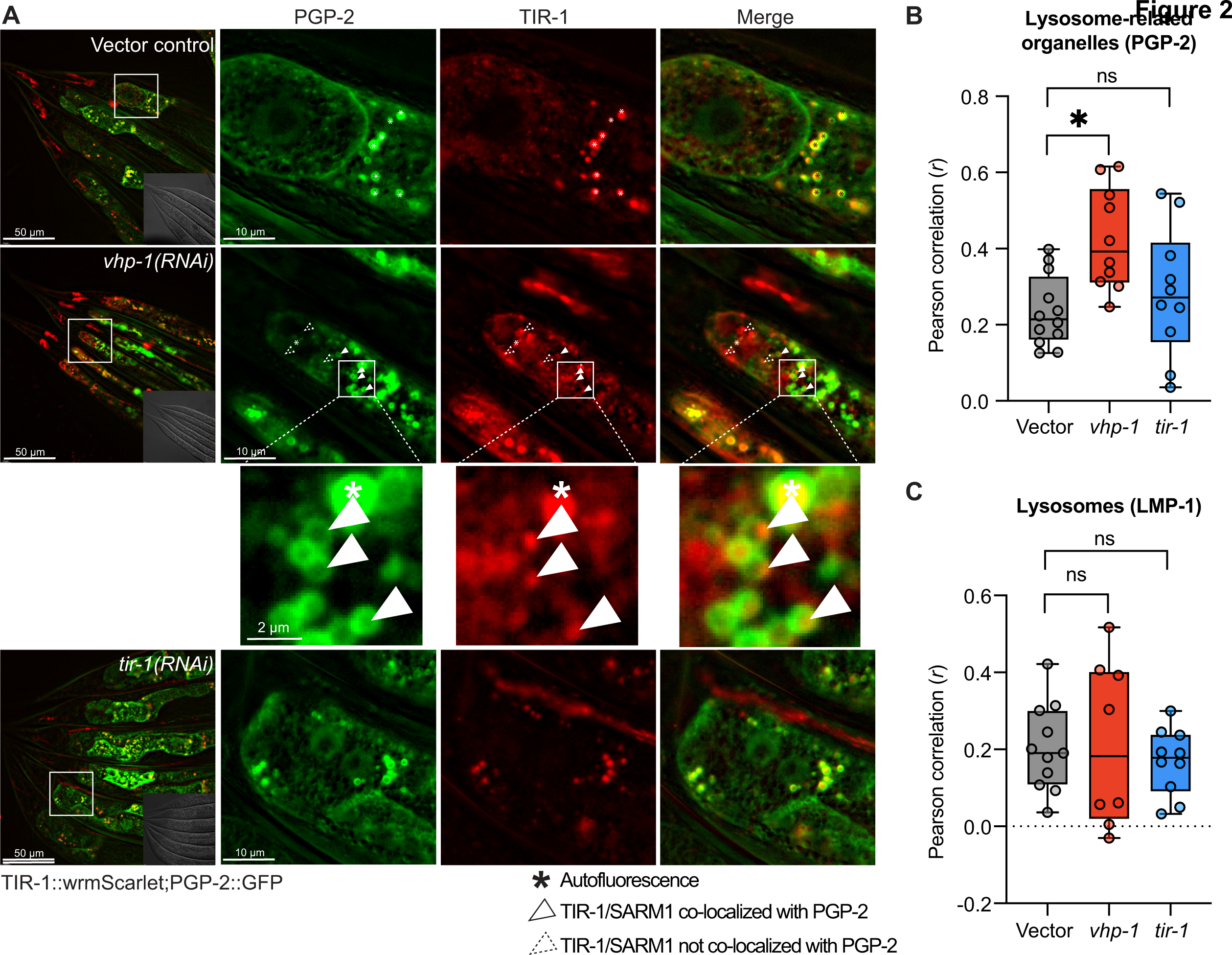
TIR-1/SARM1 co-localization with lysosome-related organelles can be visualized in live *C. elegans*. **(A)** Images of *C. elegans* TIR-1::wrmScarlet animals crossed into a PGP-2::GFP translational reporter and treated with either vector control, *vhp-1(RNAi),* or *tir-1(RNAi)*. Scale bars as indicated. Insets in the left panel represent corresponding DIC images of boxed areas. Dotted boxes indicate higher magnifications. Filled arrowheads indicate TIR-1::wrmScarlet puncta on the membranes of vesicles expressing PGP-2::GFP and dotted arrowheads indicate TIR-1::wrmScarlet puncta that do not co-localize with PGP-2. Asterisks (*) indicate autofluorescence. Quantification of TIR-1/SARM1 co-localization with **(B)** lysosome-related organelles (PGP-2) and **(C)** lysosomes (LMP-1). (B and C) Pearson correlation was calculated in each image using Fiji and the JACoP plug-in. Before analysis, non-specific autofluorescence (blue) was first subtracted from the red channel. Any remaining fluorescence was then assessed for co-localization with the green channel. Error bars represent SEM. *equals p<0.05 (one-way ANOVA with Dunnett’s multiple comparisons test). Source data for this figure is in Table S2. See also Fig. S2.

In summary, these data confirmed that TIR-1/SARM1 is expressed on lysosome-related organelles.

### A hydrophobic domain within the TIR-1/SARM1 ARM domain is required for its localization on the membranes of lysosome-related organelles and p38 PMK-1 activation

Unlike SARM1, *C. elegans* TIR-1 does not contain a putative signal peptide sequence, such as a mitochondria localizing signal.^36^ Hence, we asked if other regions of the protein were involved in its association with lysosome-related organelles. While the TIR and SAM domains of TIR-1/SARM1 are both required to activate TIR-1/SARM1 and subsequent p38 PMK-1 phosphorylation,^12^ the role of the Armadillo motif (ARM) domain in immune activation is unknown. Therefore, we asked if a region within the ARM domain of TIR-1/SARM1 was required for its association with lysosome-related organelles. Using two orthogonal *in silico* methods (DAS^37^ and Phobius^38^), we found that the ARM domain in four of the eight *C. elegans* TIR-1/SARM1 isoforms (isoform a, c, f, and g) contained a hydrophobic region (amino acids 374 to 391 in isoform a) (**Figures 3A**, **S3A, and S3B**). Using CRISPR-Cas9, we deleted the hydrophobic domain (TIR-1^ΔHD^) in the TIR-1::3xFLAG background. Co-immunostaining of anti-FLAG and anti-PGP-2 in TIR-1^ΔHD^::3xFLAG animals revealed that deleting the hydrophobic domain reduced the expression of TIR-1/SARM1 on the surface of lysosome-related organelles (**Figures 3B and 3C**). Unbiased quantification of wild-type and TIR-1^ΔHD^ mutants by the machine-learning software Aivia (Leica) revealed a significant difference in the co-localization of PGP-2 and TIR-1/SARM1 between these genotypes (**Figure 3C**). These data confirm that TIR-1/SARM1 is expressed on the membrane of lysosome-related organelles.

**Figure 3.**
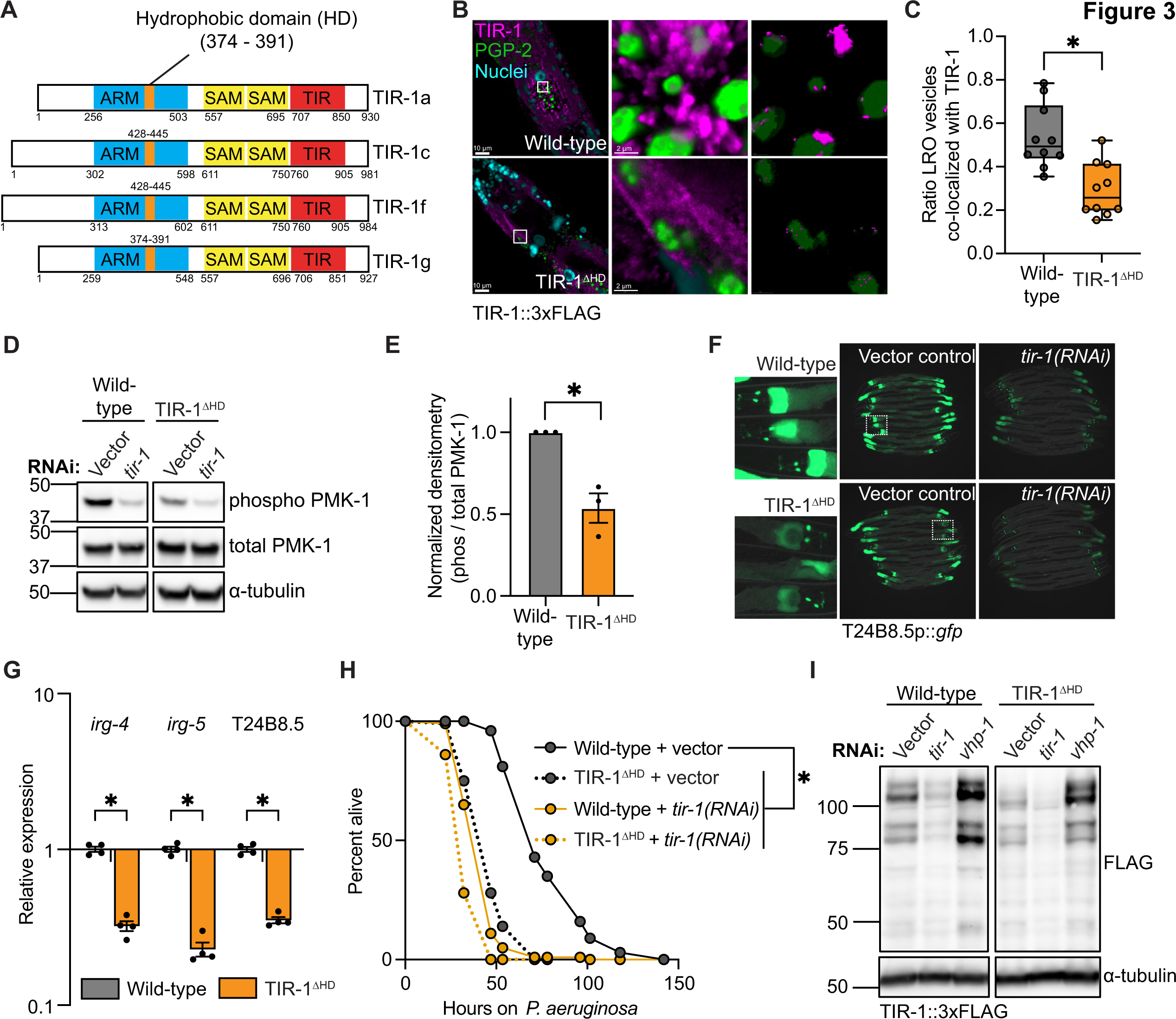
A hydrophobic domain in the TIR-1/SARM1 ARM domain is required for its localization on the membranes of lysosome-related organelles and p38 PMK-1 activation. **(A)** Schematics of the *C. elegans* TIR-1/SARM1 isoforms with a predicted hydrophobic domain as determined by Phobius^38^ and DAS.^37^ **(B)** Representative images of *C. elegans* TIR- 1::3xFLAG animals co-immunostained with both anti-FLAG and anti-PGP-2 antibodies. Images are three-dimensional reconstructions of Z-stacks. White boxes indicate higher magnifications. The last column shows only TIR-1/SARM1 (magenta) co-localized with PGP-2+ lysosome-related organelles (green). **(C)** Quantification of TIR-1/SARM1 co-localization on PGP-2+ lysosome-related organelles in wild-type and TIR-1^ΔHD^ mutants stained with both anti-FLAG and anti-PGP-2 by Aivia (Leica). *equals p<0.05 (unpaired t-test). **(D)** Representative immunoblot using anti-phospho PMK-1 and anti-total PMK-1 antibodies in wild-type and TIR-1^ΔHD^ mutants treated with vector control or *tir-1(RNAi).* **(E)** Densitometric quantification of (D) (*n*=3). Error bars represent SEM. *equals p<0.05 (unpaired t-test). **(F)** Images of wild-type and TIR-1^ΔHD^ mutants in the T24B8.5p::*gfp* transcriptional reporter background treated with either vector control or *tir-1(RNAi).* **(G)** qRT-PCR of indicated genes in wild-type and TIR-1^ΔHD^ mutants. *equals p<0.05 (two-way ANOVA with Šídák’s multiple comparisons test). **(H)** Representative *P. aeruginosa* pathogenesis assay of wild-type and TIR-1^ΔHD^ mutants treated with vector control or *tir-1(RNAi).* The difference between the wild-type and the other genotypes is significant (p<0.05, log-rank test) (n=3). Mean lifespans and statistics for all replicates are in Table S1. **(I)** Immunoblot using the anti-FLAG antibody (for TIR-1) on whole cell lysates from wild-type and TIR-1^ΔHD^ mutants within the TIR-1::3xFLAG background treated with vector control, *tir-1(RNAi)* or *vhp-1(RNAi).* Source data for this figure is in Table S2. See also Figure S3.

We also assessed whether the TIR-1/SARM1 hydrophobic domain is important for p38 PMK-1 activation. *C. elegans tir-1*^ΔHD^ mutants had significantly reduced levels of active p38 PMK-1 at baseline (*e.g*., in the absence of infection) (**Figures 3D and 3E**). In addition, baseline expression of the p38 PMK-1-dependent transcriptional immune reporter T24B8.5p::*gfp* was subtly suppressed in *C. elegans tir-1*^ΔHD^ mutants, most noticeably in the anterior intestine (**Figure 3F**). Accordingly, *tir-1*^ΔHD^ mutants also expressed significantly lower transcription of the p38 PMK-1-dependent targets *irg-4, irg-5,* and T24B8.5 by qRT-PCR (**Figure 3G**). Importantly, *C. elegans tir-1*^ΔHD^ mutants were also more susceptible to killing by *P. aeruginosa* than wild-type animals (**Figure 3H**). These data demonstrated that the hydrophobic domain of TIR-1/SARM1 is required for p38 PMK-1 activation and pathogen resistance. Intriguingly, the susceptibility of *tir-1*^ΔHD^ to *P. aeruginosa* was similar to that of *tir-1(RNAi)* animals (**Figure 3H**), despite the relatively small decrease in p38 PMK-1 phosphorylation (**Figure 3D-E**) and T24B8.5p::*gfp* (**Figure 3F**) expression in *tir-1*^ΔHD^ mutants. Thus, these data suggest that the *tir-1*^ΔHD^ mutation affects *C. elegans* susceptibility to bacterial infection predominantly by blocking a pathogen-inducible host response, rather than by affecting the basal expression of immune effectors. We explore this hypothesis further below.

We confirmed that TIR-1/SARM1 protein was transcribed in *tir-1*^ΔHD^ mutants at the same levels as wild-type animals (**Figure S3C**). Using an anti-FLAG antibody, we also quantified TIR-1::3xFLAG and TIR-1^ΔHD^::3xFLAG protein by immunoblot. All isoforms were expressed in TIR-1^ΔHD^ mutants, albeit at subtly lower levels (**Figures 3I and S3D**). Importantly, knockdown of *vhp-1*, which engaged feedforward transcription and translation of *tir-1* as described in the companion manuscript, robustly induced TIR-1^ΔHD^::3xFLAG protein to similar levels as wild-type TIR-1::3xFLAG protein (**Figure 3I**). In addition, knockdown of *tir-1* further suppressed TIR-1/SARM1 protein expression in the TIR-1^ΔHD^ mutant (**Figure 3I**). Thus, the TIR-1^ΔHD^::3xFLAG protein is appropriately translated in *tir-1*^ΔHD^ mutants.

Cryo-electron microscopic studies of *C. elegans* TIR-1/SARM1 show that the hydrophobic domain is located within a pocket of the ARM domain of the protein (**Figure S3E**).^39^ Thus, in addition to the possibility that the hydrophobic domain directly facilitates the association of TIR-1/SARM1 with the membrane of lysosome-related organelles, it is also possible the TIR-1^ΔHD^ mutation de-stabilizes the ARM domain or full length TIR-1/SARM1 protein in a manner that prevents its interaction with the membrane of lysosome-related organelles. In either case, these studies of the TIR-1^ΔHD^ mutant provide important confirmation that TIR-1/SARM1 is expressed on the membranes of lysosome-related organelles.

### The pathogen-derived virulence effector pyocyanin alkalinizes and condenses lysosome-related organelles in intestinal epithelial cells

To determine how pathogen infection affects lysosome-related organelles in *C. elegans*, we used LysoTracker Red, a dye that marks acidic organelles (including lysosome-related organelles),^35,40^ to stain animals that were infected with strains of *P. aeruginosa* with differing virulence potentials (**Figure 4A**). We observed that the intensity of LysoTracker Red staining was reduced in animals infected with virulent strains of *P. aeruginosa,* but not in the less virulent *P. aeruginosa* PAK (**Figures 4A and 4B**). Of note, *P. aeruginosa-*mediated depletion of LysoTracker Red was not dependent on *pmk-1* or *tir-1* (**Figure S4A**). These data indicate that depletion of lysosome-related organelles during pathogen infection does not occur downstream of p38 PMK-1 (*i.e.*, as a consequence of immune activation), but rather may be an initiating event of innate immune sensing and p38 PMK-1 activation.

**Figure 4.**
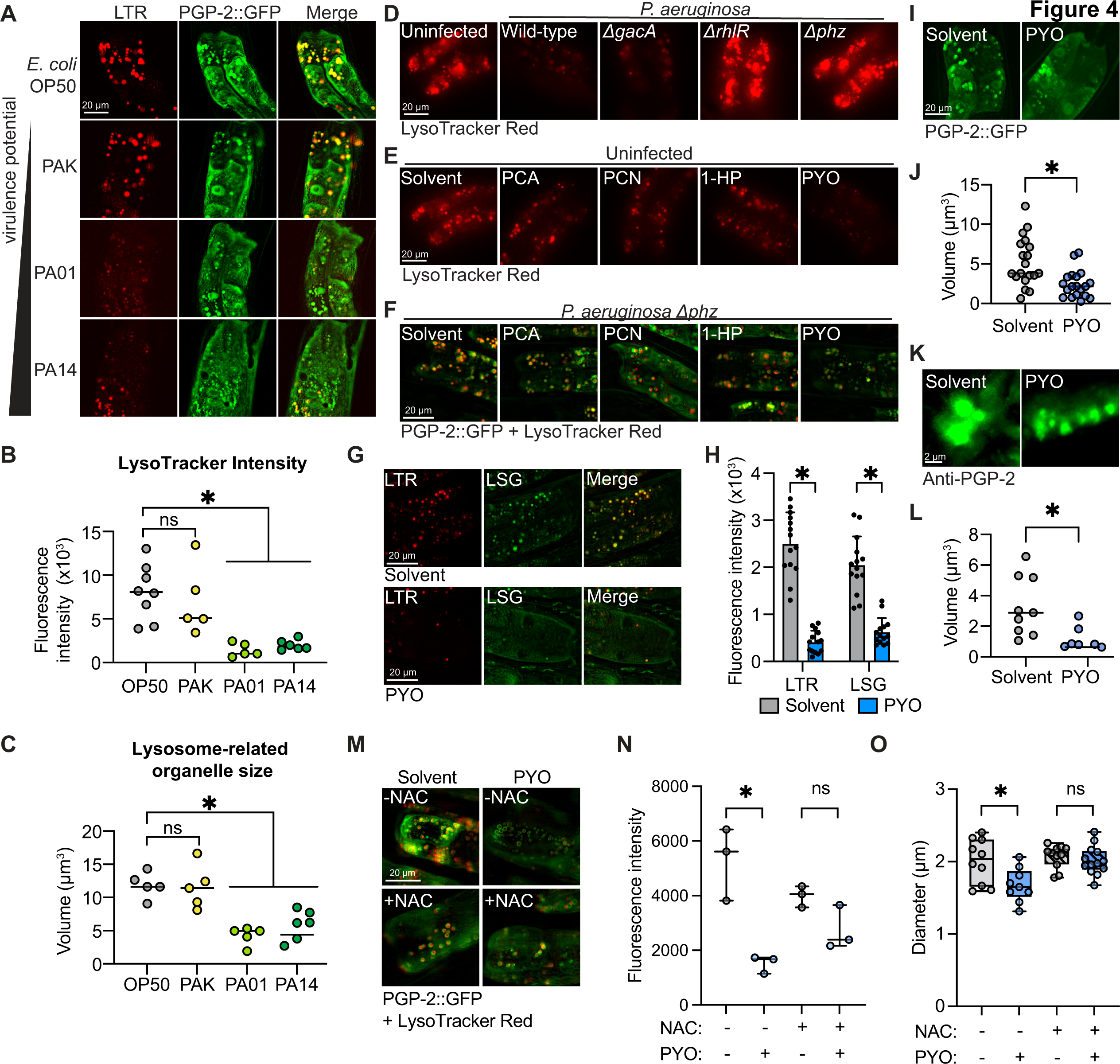
The pathogen-derived virulence effector pyocyanin alkalinizes and condenses lysosome-related organelles in intestinal epithelial cells. **(A)** *C. elegans* PGP-2::GFP translational reporters were stained with LysoTracker Red (LTR, 1 µM) and transferred to the indicated *P. aeruginosa* strains for 4-6 hrs. **(B)** Quantification of LysoTracker Red intensity in (A) using the LasX analysis software (Leica). *equals p<0.05 (one-way ANOVA with Dunnett’s multiple comparisons test). **(C)** Quantification of PGP-2+ vesicle size using the LasX three-dimensional analysis software (Leica). *equals p<0.05 (one-way ANOVA with Dunnett’s multiple comparisons test). **(D)** Images of wild-type LysoTracker Red-stained *C. elegans* uninfected or infected with indicated *P. aeruginosa* mutants for 4 hrs. **(E)** Images of uninfected wild-type LysoTracker Red-stained animals supplemented with PCA (200 µM), PCN (200 µM), 1-HP (20 µM) or PYO (200 µM) for 4 hrs. **(F)** Images of *P. aeruginosa Δphz-*infected wild-type PGP-2::GFP animals stained with LysoTracker Red in presence or absence phenazine supplementation (same concentrations as (E)). **(G)** Images of wild-type animals treated with solvent control or PYO (200 µM) and stained with both LysoTracker Red (LTR, 1 µM) and LysoSensor Green (LSG, 1 µM). **(H)** Quantification of fluorescence intensity in (G). *equals p<0.05 (two-way ANOVA with Šídák’s multiple comparisons test). **(I-J)** Wild-type *C. elegans* PGP-2::GFP animals treated with solvent or PYO (200 µM) for 4 hrs. (I) Representative images and (J) quantification of PGP-2::GFP volume. *equals p<0.05 (unpaired t-test). **(K-L)** Wild-type *C. elegans* treated with solvent or PYO (200 µM) for 4 hrs and immunostained with the anti-PGP-2 antibody. (K) Representative images and (L) quantification of PGP-2+ vesicle volume. *equals p<0.05 (unpaired t-test). Scale bars as indicated. **(M)** Images of wild-type PGP-2::GFP animals pre-treated with solvent control or N-acetylcysteine (NAC, 5 mM) for 2 hrs followed by exposure to either solvent control or PYO (200 µM) for 4 hrs. (**N**) Quantification of LysoTracker Red fluorescence intensity in (M). *equals p<0.05 (one-way ANOVA with Tukey’s multiple comparisons test). (**O**) Quantification of diameter of PGP-2::GFP(+) vesicles in (M). *equals p<0.05 (one-way ANOVA with Tukey’s multiple comparisons test). Source data for this figure is in Table S2. See also Figure S4.

In conjunction, we assessed *C. elegans* lysosome-related organelles using the PGP-2::GFP translational reporter (**Figures 4A and 4C**). The volume of PGP-2::GFP+ vesicles was significantly reduced during pathogen infection in a manner dependent on the virulence potential of the bacterial pathogen (**Figure 4C**). As previously reported,^35,40^ we also observed that LysoTracker Red+ vesicles co-localized with PGP-2::GFP (**Figure 4A**). Similar to the observations with LysoTracker Red (**Figure S4A**), the reduction in size of PGP-2::GFP+ vesicles during pathogen infection was not dependent on *pmk-1* or *tir-1* (**Figure S4B**). In summary, *P. aeruginosa* infection alkalinizes (**Figures 4A and 4B**) and condenses (**Figures 4A and 4C**) lysosome-related organelles.

To determine if *P. aeruginosa* specifically targets lysosome-related organelles in *C. elegans* during infection, we assessed mutants with deletions in master regulators of virulence. LysoTracker Red staining of lysosome-related organelles was still markedly reduced in *P. aeruginosa ΔgacA* mutants, but not upon exposure to *P. aeruginosa ΔrhlR* mutants (**Figure 4D**). These data suggest that *rhlR* regulates a gene program in *P. aeruginosa* that reduces the size of host lysosome-related organelles. Thus, we screened transposon mutants for 155 *rhlR-* dependent genes to identify individual pseudomonal virulence effectors that target lysosome-related organelle integrity.^41^ Four of the 155 mutants screened phenocopied the *ΔrhlR* mutant with respect to LysoTracker Red staining of *C. elegans* lysosome-related organelles, two of which were mutations in *phzA2* and *phzB2* (**Table S3**). Intriguingly, each of these genes is involved in the production of phenazines, toxic metabolites that are secreted by *P. aeruginosa*. Accordingly, a *P. aeruginosa* strain with a clean deletion in the phenazine biosynthetic operon (*Δphz*) was unable to reduce the size and number of lysosome-related organelles during infection (**Figure 4D**).

To determine if a single phenazine metabolite caused the effects of *P. aeruginosa* on lysosome-related organelles, we treated animals with phenazine-1-carboxylic acid (PCA), phenazine-1-carboxamide (PCN), 1-hydroxyphenazine (1-HP), and pyocyanin (**Figure 4E**). Of the four phenazine metabolites, treatment with only pyocyanin fully recapitulated the pattern of LysoTracker Red staining observed during infection with wild-type *P. aeruginosa* (**Figure 4E**). In addition, treatment with only pyocyanin, and not the other phenazines, restored the suppression of PGP-2+, LysoTracker Red+ lysosome-related organelles in *C. elegans* exposed to the *P. aeruginosa Δphz* mutant (**Figure 4F**). These data suggested that the secreted virulence effector pyocyanin disrupts the pH of lysosome-related organelles in the *C. elegans* intestine. To confirm this observation, we co-stained pyocyanin-treated and control animals with both LysoTracker Red and LysoSensor Green, a dye with a pKa of ∼5.2, which is more sensitive to changes in pH. Animals treated with pyocyanin had significantly less staining of both LysoTracker Red and LysoSensor Green (**Figures 4G and 4H**). Importantly, pyocyanin did not change the pH of the media on which animals were grown (**Figure S4C**). These data demonstrated that pyocyanin alkalinizes lysosome-related organelles.

We then used two orthogonal approaches to determine if pyocyanin treatment recapitulated the change in lysosome-related organelle size we observed upon *P. aeruginosa* infection. Treatment with pyocyanin significantly reduced the volume of PGP-2::GFP+ vesicles (**Figures 4I and 4J**). In addition, immunostaining of lysosome-related organelles with an anti-PGP-2 antibody also revealed that pyocyanin treatment led to a reduction in size of lysosome-related organelles (**Figures 4K and 4L**).

The secreted virulence effector pyocyanin is a redox active molecule that induces oxidative stress in host cells during *P. aeruginosa* infection.^42–44^ We hypothesized that the alkalinization and condensation of lysosome-related organelles were caused by oxidative stress induced by pyocyanin. Consistent with this hypothesis, pre-treatment of *C. elegans* with the anti-oxidant N-acetylcysteine (NAC) prevented the suppression of LysoTracker Red staining caused by pyocyanin (**Figures 4M and 4N**). In addition, NAC treatment abrogated the pyocyanin-induced reduction in size of PGP-2::GFP+ lysosome-related organelles (**Figures 4M and 4O**).

In summary, oxidative stress induced by pathogen-derived pyocyanin, a redox active virulence effector, alkalinized and condensed lysosome-related organelles in *C. elegans* intestinal epithelial cells.

### Pyocyanin triggers aggregation of TIR-1/SARM1 on lysosome-related organelles to activate p38 PMK-1 intestinal immunity

We previously found that TIR-1/SARM1 multimerizes in response to physical crowding, which activates its intrinsic NAD^+^ hydrolase activity.^12^ In addition, we observed that TIR-1/SARM1 multimerization is augmented with increasing pH.^12^ We therefore hypothesized that the pyocyanin-induced condensation and alkalinization of lysosome-related organelles triggers aggregation of TIR-1/SARM1, which activates the p38 PMK-1 immune pathway. Supporting this model of immune activation in *C. elegans*, treatment with pyocyanin, but not the other phenazines, activated p38 PMK-1 phosphorylation in wild-type animals (**Figures 5A and 5B**). In addition, only pyocyanin robustly induced TIR-1::3xFLAG protein in wild-type animals (**Figures S5A and S5B**), consistent with our data demonstrating that p38 PMK-1 phosphorylation drives induction of TIR-1/SARM1 protein.

**Figure 5.**
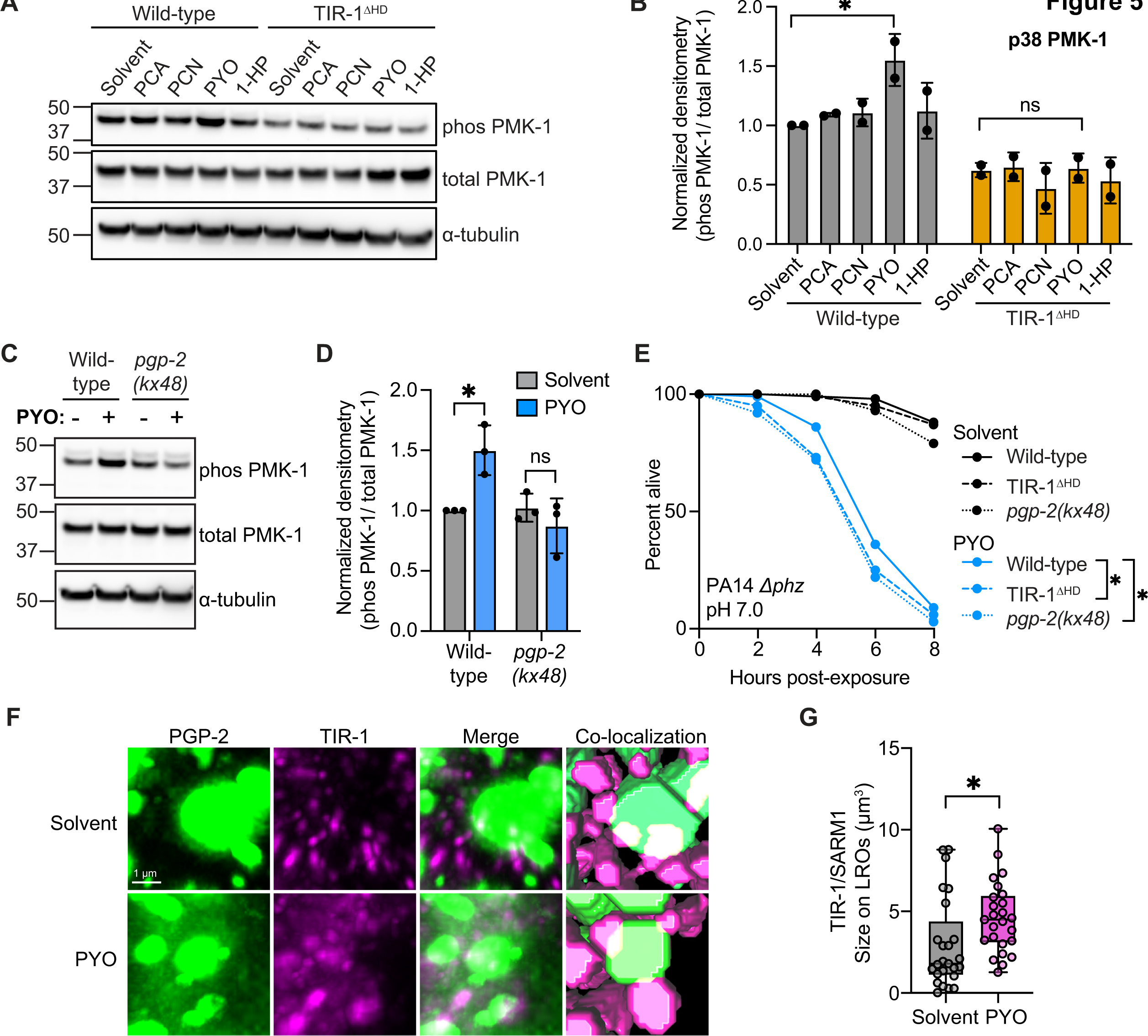
Pyocyanin triggers aggregation of TIR-1/SARM1 on lysosome-related organelles and activates p38 PMK-1 intestinal immunity. **(A)** Representative immunoblot of whole cell lysates isolated from wild-type and TIR-1^ΔHD^ mutants treated with the indicated phenazines for 4-6 hrs and probed with anti-phospho PMK-1, anti-total PMK-1, and anti-⍺-tubulin antibodies. **(B)** Densitometric quantification of conditions in (A) (*n*=2). *equals p<0.05 (two-way ANOVA with Tukey’s multiple comparisons test). **(C)** Representative immunoblot of whole cell lysates isolated from wild-type and TIR-1^ΔHD^ mutants in the TIR-1::3xFLAG background treated with indicated phenazines for 4-6 hrs and probed with anti-FLAG and anti-⍺-tubulin antibodies. **(D)** Densitometric quantification of conditions in (C) (*n=*3). *equals p<0.05 (two-way ANOVA with Tukey’s multiple comparisons test). **(E)** Representative pyocyanin toxicity “fast-kill” assay of wild-type, TIR-1^ΔHD^, and *pgp-2(kx48)* mutants. The difference between the wild-type and the other genotypes is significant (p<0.05, log-rank test) (n=3). **(F)** Representative images of animals treated with solvent control or PYO (200 µM) co-immunostained with anti-FLAG (for TIR-1) and anti-PGP-2. Last column represents Aivia (Leica) render of TIR-1/SARM1 and PGP-2 co-localization. (**G**) Quantification of TIR-1/SARM1 puncta size on PGP-2+ vesicles in co-immunostained animals in the presence or absence of PYO. *equals p<0.05 (unpaired t-test). All phenazines were used at 200 µM except for 1-HP (20 µM). Scale bars as indicated. Mean lifespans and statistics for all replicates are in Table S1. Source data for this figure is in Table S2. See also Figure S5.

We previously discovered that the phenazine PCN is sensed by the *C. elegans* nuclear hormone receptor NHR-86, a ligand-gated transcription factor.^20^ When activated, NHR-86 directly induces an anti-pathogen transcriptional program and provides protection against killing by *P. aeruginosa* independent of the p38 PMK-1 immune pathway.^20,45^ Consistent with these data, PCN did not increase p38 PMK-1 phosphorylation (**Figures 5A and 5B**) or TIR-1::3xFLAG protein (**Figures S5A and S5B**).

Importantly, the hydrophobic domain of TIR-1/SARM1, which was required for its localization to lysosome-related organelles (**Figure 3**), was also necessary for the activation of p38 PMK-1 by pyocyanin (**Figures 5A and 5B**). In addition, the *tir-1*^ΔHD^ mutation abrogated the pyocyanin-induced increase in TIR-1/SARM1 translation observed in wild-type animals (**Figures S5A and S5B**).

Importantly, activation of p38 PMK-1 by pyocyanin treatment requires the presence of PGP-2+ lysosome-related organelles. Ablation of this cellular compartment in *pgp-2(kx48)* loss-of-function mutants suppressed the pyocyanin-mediated activation of p38 PMK-1 (**Figures 5C and 5D**). *C. elegans pgp-2(kx48)* mutants also had significantly higher transcription of *tir-1* mRNA compared to wild-type animals (**Figure S3C**), suggesting that a compensatory response is engaged in this genetic background to increase *tir-1* transcription. Together, these data demonstrate that alkalinization and contraction of lysosome-related organelles by the pathogen-derived toxic metabolite pyocyanin is an upstream event that activates p38 PMK-1 innate immune signaling.

We assessed the contribution of lysosome-related organelles toward resistance to pyocyanin. Exposure to pyocyanin at an elevated pH led to rapid killing of wild-type *C. elegans* (**Figure 5E**), consistent with previous studies.^46^ Importantly, both *C. elegans pgp-2(kx48)* and *tir-1*^ΔHD^ mutants were hypersusceptible to pyocyanin-mediated toxicity in this pathogenesis assay, which is also called the “fast kill” assay (**Figure 5E**). *C. elegans pgp-2(kx48)* mutants were also hypersusceptible to killing by *C. elegans* in an intestinal infection assay (the “slow-kill” assay) (**Figure S5C**). Thus, these results indicate that intact lysosome-related organelles, as well as TIR-1/SARM1 co-localization on these vesicles, are important for resistance against the pathogen-derived virulence effector pyocyanin.

We considered a model where pyocyanin treatment liberates TIR-1/SARM1 from lysosome-related organelles to form cytoplasmic puncta and activate the p38 PMK-1 pathway. We co-immunostained PGP-2 and TIR-1/SARM1 in animals treated with pyocyanin or solvent control. Pyocyanin treatment did not significantly change overall TIR-1/SARM1 localization to the membranes of lysosome-related organelles (**Figure S5D**), suggesting that TIR-1/SARM1 is not released from the vesicles under these conditions. We then asked whether pyocyanin treatment led to increased TIR-1/SARM1 puncta formation on lysosome-related organelles. Using Aivia (Leica), we quantified the size (volume) of TIR-1/SARM1 puncta on lysosome-related organelles. Treatment with pyocyanin led to a significant increase in TIR-1/SARM1 puncta size on lysosome-related organelles compared to solvent control alone (**Figures 5F and 5G**).

In summary, pyocyanin is a virulence effector produced by *P. aeruginosa* that triggers alkalinization and contraction of lysosome-related organelles, aggregation of TIR-1/SARM1 on these organelles, and activation of p38 PMK-1 intestinal immunity (**Figure 6**).

**Figure 6.**
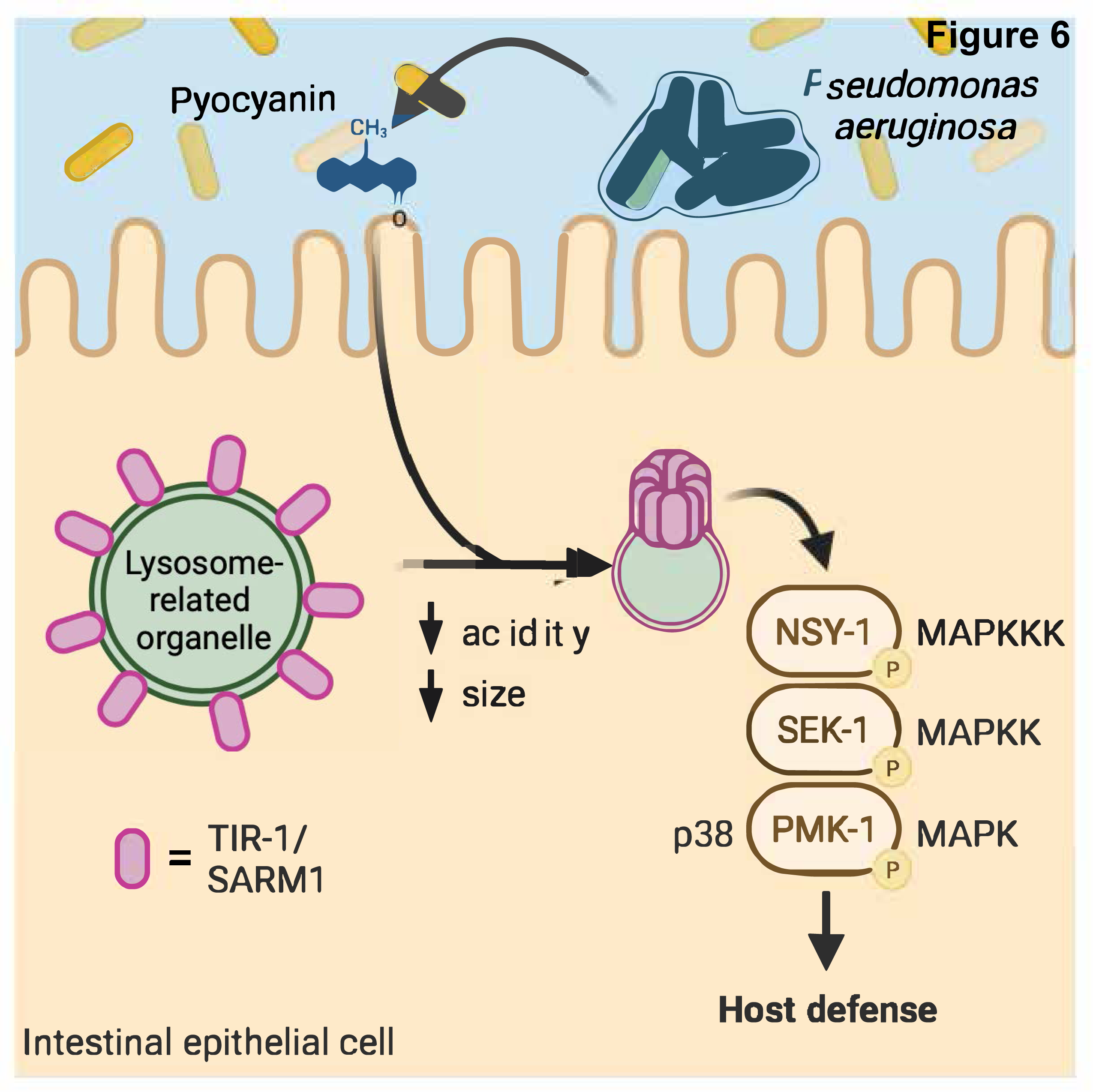
Pathogen effector-triggered aggregation of TIR-1/SARM1 on lysosome-related organelles activates intestinal immunity.

## DISCUSSION

In this study, we characterized a mechanism of enzymatic TIR activation in animal immunity. This family of proteins is strongly conserved in evolution, with essential roles in plant immunity and in bacterial defense against phage infection.^5^ However, it was not previously known how these proteins function in pathogen sensing in animals. We showed that the single NADase protein with a TIR domain in *C. elegans* is expressed on the membranes of a specific subset of lysosomes, called lysosome-related organelles. Pyocyanin, a toxic phenazine secreted by *P. aeruginosa*, induces oxidative stress, which alkalinized and condensed lysosome-related organelles causing TIR-1/SARM1 to form higher-order multimers on the membranes of these organelles. We previously demonstrated that TIR-1/SARM1 multimerization engages its intrinsic NAD^+^ hydrolase activity to activate the p38 PMK-1 immune pathway.^12^ Accordingly, activation p38 PMK-1 signaling by pyocyanin requires TIR-1/SARM1 expression on lysosome-related organelles.

In a companion study co-submitted with this manuscript, we discovered that maintenance of lysosomal integrity restrains deleterious propagation of p38 PMK-1 innate immune signaling. From a forward genetic screen for new immune regulators, we identified a previously uncharacterized gene that we named *rotr-1* and discovered that this gene is necessary to maintain the size of lysosome-related organelles. Accordingly, condensation of lysosome-related organelles in *rotr-1* mutants promoted TIR-1/SARM1 aggregation and constitutive hyperactivation of p38 PMK-1 immune pathway. Thus, a forward genetic screen identified *C. elegans* mutants that recapitulated the pathogen-induced damage to the specific lysosomal compartment that expressed TIR-1/SARM1. These data provide an orthologous confirmation for the cell biological characterization of p38 immune activation in this manuscript, and demonstrate that the lysosome – TIR-1/SARM1 – p38 axis is also essential for immune homeostasis in the animal intestine.

In addition to characterizing a TIR enzyme in animal innate immunity, our data uncover a prominent role for lysosome-related organelles in pathogen sensing and immune activation. These findings provide a key insight into the biological function of this prominent, yet incompletely understood, subcellular compartment that is present in all metazoans. Lysosome-related organelles are terminal endocytic vesicles that function as sites for storage and secretion of a diverse array of substances in all metazoan animals.^47^ In humans, specialized lysosome-related organelles within leukocytes, for example, are reservoirs for toxic effector molecules.^48^ In *C. elegans,* lysosome-related organelles, also called gut granules, are ubiquitous throughout intestinal tissues. These vesicles are acidic compartments that store zinc and heme,^33,35,49–51^ are involved in ascaroside biosynthesis,^52^ and have been implicated in stress resistance. Loss of lysosome-related organelles inhibited transcriptional activation of some putative immune genes.^53^ Here, we showed that lysosome-related organelles are scaffolds for an innate immune signaling regulator that change dynamically during infection to enable sensing of a pathogen-derived toxin. These data suggest that the evolutionarily ancient function of lysosome-related organelles may be in stress sensing, offering a clue to their function in higher order hosts.

Finally, our findings link surveillance of a bacterial pattern of pathogenesis to activation of p38 PMK-1 signaling, solving a 20-year-old mystery of how this core innate immune pathway is activated in *C. elegans* during pathogen infection. Nematodes do not utilize canonical mechanisms of pattern recognition to identify bacterial pathogens and activate immune signaling pathways, as nematodes lost traditional pattern recognition receptors in evolution. Instead, *C. elegans* evolved mechanisms to monitor for the effects of pathogen infection on the host to identify pathogen attack. Prior to this study, it was not previously known if surveillance of “patterns of pathogenesis” link sensing of bacterial infection to the activation of protective immune signaling cascades in the *C. elegans* intestinal epithelium.

Phenazines are toxic metabolites made exclusively by *Pseudomonas* species.^54^ These molecules are produced only when a community of pseudomonal bacteria has grown to a specific density, under the control of so-called quorum sensing mechanisms, and play particularly important roles in bacterial physiology.^55,56^ For example, phenazines promote aerobic respiration within the relatively anoxic interior of a pseudomonal biofilm by serving as electron shuttles.^57^ During infection, these molecules are *bona fide* virulence effectors that perturb mitochondrial function.^26,58^ We previously found that one phenazine, PCN, is sensed directly in *C. elegans* as a readout of *P. aeruginosa* virulence potential.^20^ Here we demonstrate that *C. elegans* sense the toxic effects of another phenazine, pyocyanin, by surveilling the state of lysosome-related organelles to initiate protective innate immune defenses in *C. elegans*. These findings collectively underscore the importance of phenazines to pseudomonal virulence and demonstrate that *C. elegans* evolved specific mechanisms to detect these metabolites, which are essential for bacterial pathogenesis.

In conclusion, our results establish the mechanism by which the effects of a toxin produced by *P. aeruginosa* are detected by *C. elegans* to activate innate immunity. Our study defines the mechanism of enzymatic TIR activation in animal immunity and demonstrates that lysosome-related organelles are primordial immune organelles, uniquely positioned within intestinal epithelia to recognize and respond to the effects of a virulent pathogen on the host.

### Limitations of Current Study

We observed two seemingly distinct populations of TIR-1/SARM1 puncta: those associated with lysosome-related organelles and those in the cytoplasm. The TIR-1/SARM1 puncta on the membrane of lysosome-related organelles dramatically increased in size following exposure to pyocyanin. We hypothesize that the cytoplasmic TIR-1/SARM1 puncta, which did not seem to be associated with lysosome-related organelles, are membrane-free, higher ordered protein assembles. In a companion study, we discovered that activation of the p38 PMK-1 pathway leads to the feedforward propagation of immune signaling by increase the transcription, translation and multimerization of TIR-1/SARM. Thus, the TIR-1/SARM1 cytoplasmic puncta may formed as a consequence of the feedforward propagation of *tir-1* transcription by the p38 PMK-1 immune pathway.

## ACKNOWLEDGEMENTS

The authors thank Greg Hermann for the anti-PGP-2 antibody, Alexander Soukas for *C. elegans* strains and helpful discussions, Kit-Yi Yam for helpful suggestions regarding the image analyses, and Melanie Trombly and Milka Kostic for critical reading of the manuscript. The research was supported by R01 AI130289 (to R.P.W.), R01 AI159159 (to R.P.W.), R21 AI163430 (to R.P.W.), F30 DK127690 (to S.Y.T.), T32 AI095213 (to S.Y.T.), F30 AI150127 (to N.D.P.), T32 AI132152 (to N.D.P.), and T32 GM107000 (to S.Y.T. and N.D.P.). Some strains were provided by the *Caenorhabditis* Genetics Center, which is funded by the NIH Office of Research Infrastructure Programs (P40 OD010440). The funders had no role in study design, data collection and analysis, decision to publish, or preparation of the manuscript.

## AUTHOR CONTRIBUTIONS

Conceptualization: SYT, RPW; Methodology: SYT, RPW; Investigation: SYT, KAW, AP; Visualization: SYT; Funding acquisition: RPW; Project administration: RPW; Supervision: RPW; Writing – original draft, SYT, RPW; Writing – review & editing, SYT, RPW

## INCLUSION AND DIVERSITY STATEMENT

We support inclusive, diverse, and equitable conduct of research.

## STAR METHODS

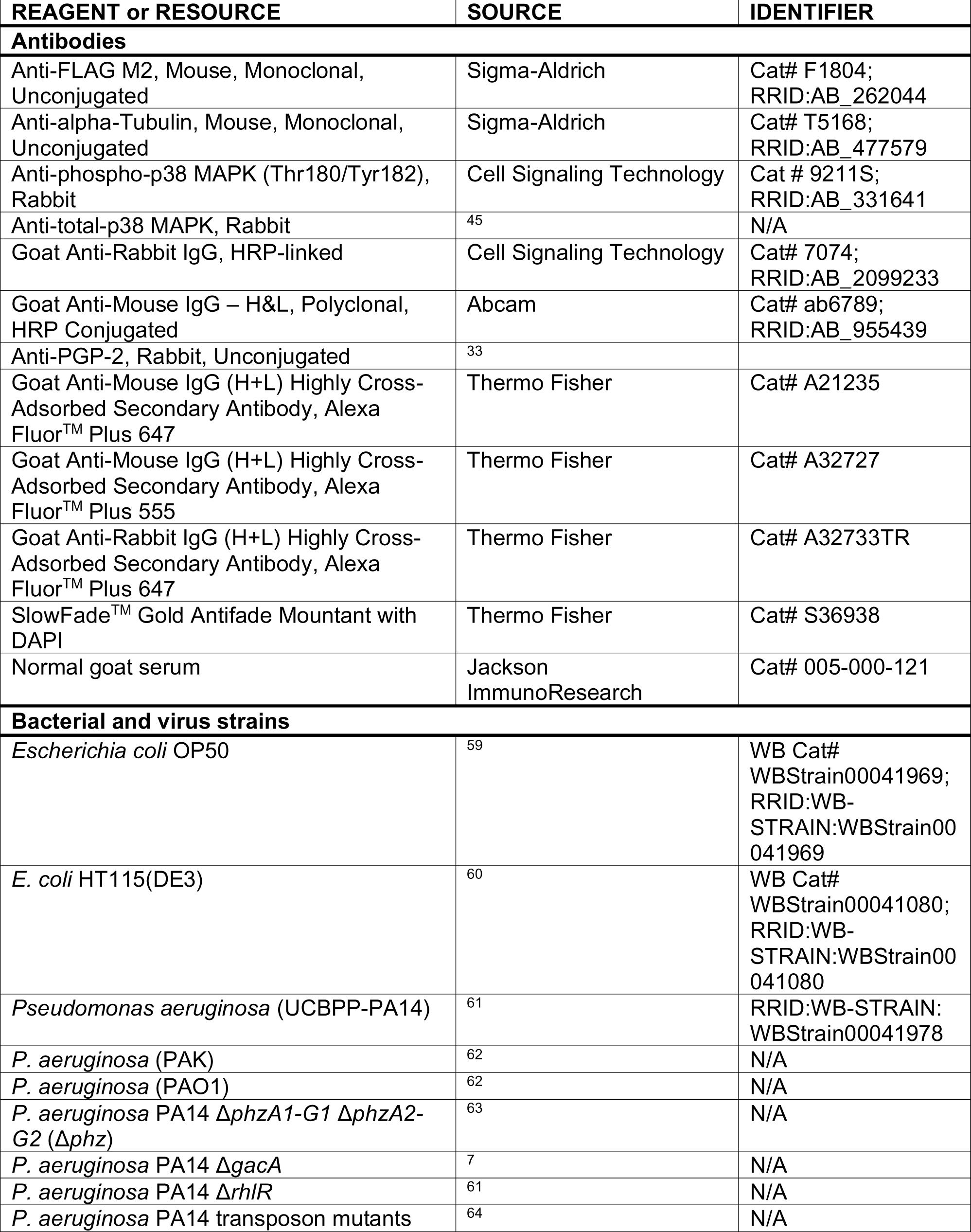
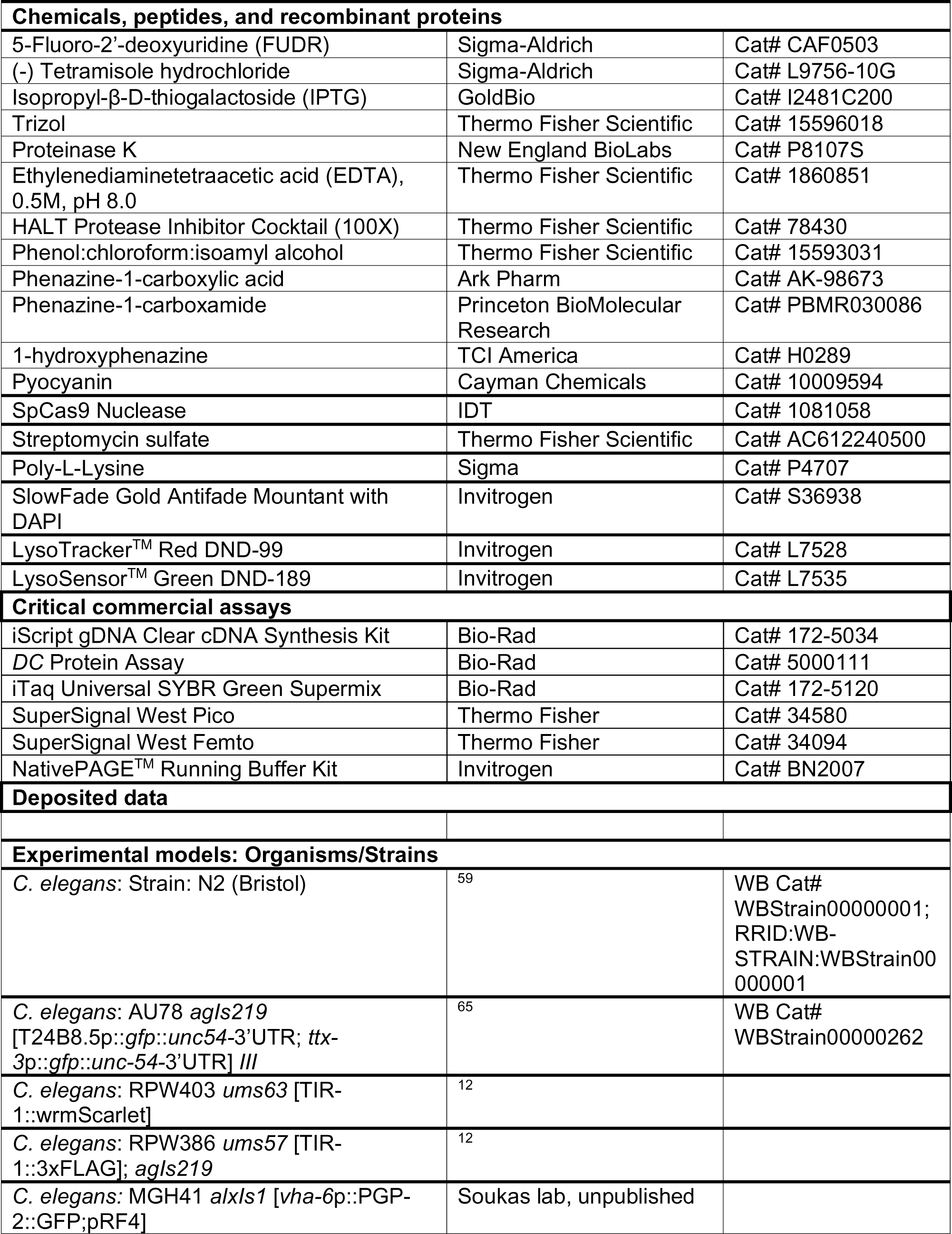
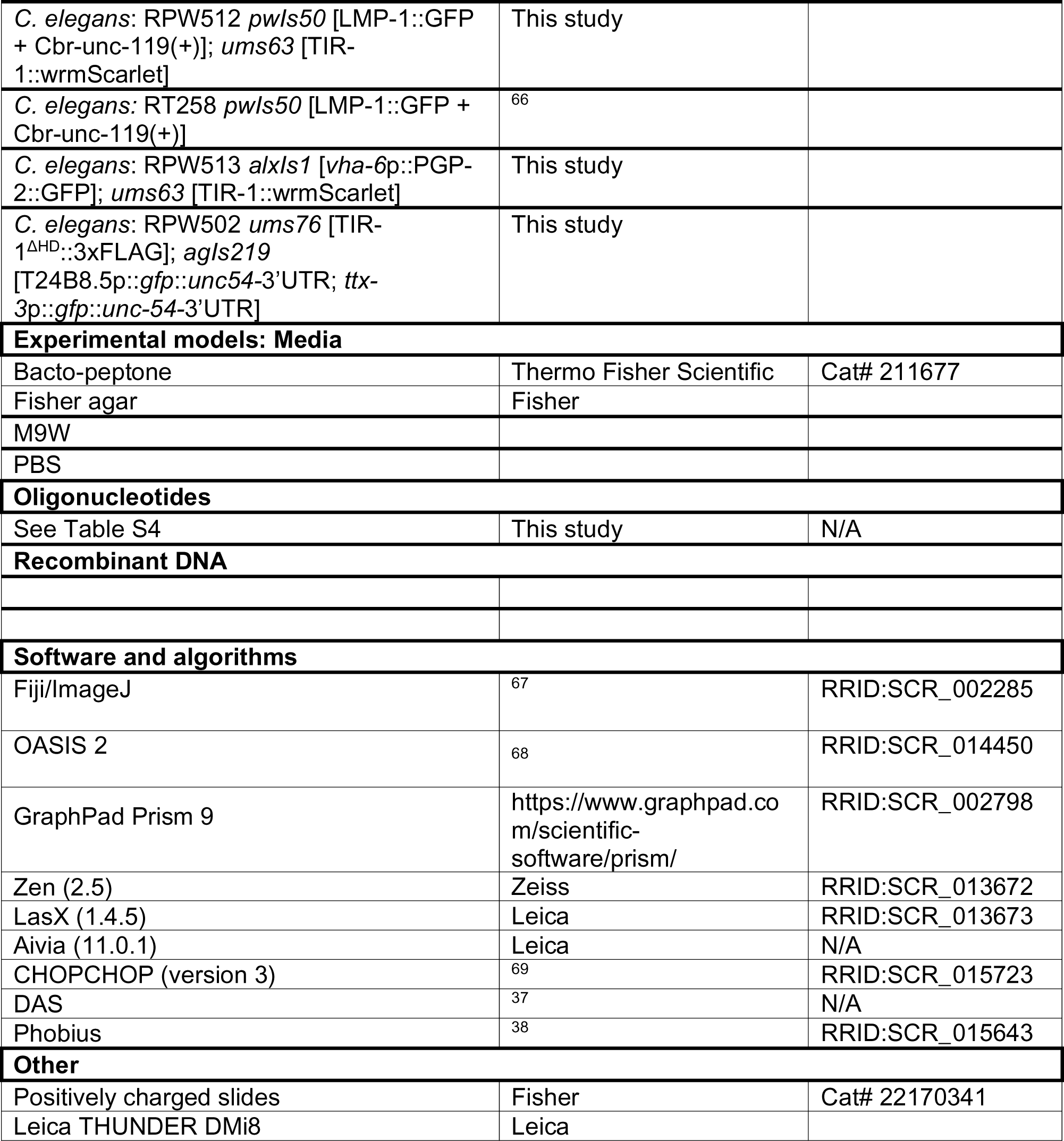
KEY RESOURCES TABLE

## RESOURCE AVAILABILITY

### Lead contact

Further information and requests for resources or reagents should be directed to and will be fulfilled by the lead contact, Read Pukkila-Worley.

### Material availability

Strains and reagents generated in this study are available upon request.

### Data and code availability

This paper does not report original code.

## EXPERIMENTAL MODEL AND SUBJECT DETAILS

### C. elegans strains

The previously published *C. elegans* strains used in this study were: N2 Bristol,^59^ AU78 *agIs219* [T24B8.5p::*gfp*::*unc54-*3’UTR; *ttx-3*p::*gfp*::*unc-54-*3’UTR] *III,*^65^ RPW403 *ums63* [TIR-1::wrmScarlet],^12^ RPW386 *ums57* [TIR-1::3xFLAG]; *agIs219* [T24B8.5p::*gfp*::*unc54-*3’UTR; *ttx-3*p::*gfp*::*unc-54-*3’UTR] *III,*^12^ MGH41 *alxIs1* [*vha-6*p::PGP-2::GFP;pRF4] (unpublished), RT258 *pwIs50* [LMP-1::GFP + Cbr-unc-119(+)].^66^ The strains developed in this study were: RPW512 *pwIs50* [LMP-1::GFP + Cbr-unc-119(+)]; *ums63* [TIR-1::wrmScarlet], RPW513 *alxIs1* [*vha-6*p::PGP-2::GFP]; *ums63* [TIR-1::wrmScarlet], RPW502 *ums76* [TIR-1^ΔHD^::3xFLAG]; *agIs219* [T24B8.5p::*gfp*::*unc54-*3’UTR; *ttx-3*p::*gfp*::*unc-54-*3’UTR].

### *C. elegans* growth conditions

*C. elegans* strains were maintained on standard nematode growth medium (NGM) plates [0.25% Bacto-peptone, 0.3% sodium chloride, 1.7% agar (Fisher agar), 5 μg/mL cholesterol, 25 mM potassium phosphate pH 6.0, 1 mM magnesium sulfate, 1 mM calcium chloride] with *E. coli* OP50 as a food source, as described.^59^

### Bacterial strains

Bacteria used in this study were *Escherichia coli* (*E. coli*) OP50,^59^ *E. coli* HT115(DE3),^60^ and *Pseudomonas aeruginosa* strains PA14,^61^ PAO1,^62^ PAK,^62^ PA14 Δ*phzA1-G1* Δ*phzA2-G2* (Δ*phz*),^63^ PA14 Δ*gacA,*^7^ PA14 Δ*rhlR,*^61^ and PA14 transposon mutants.^64^

### Bacterial growth conditions

*E. coli* OP50 were grown in LB broth supplemented with 0.175 mg/mL streptomycin at 37 °C for 16-18 hrs at 250 rpm. *P. aeruginosa* strains were grown in LB broth at 37 °C for 14-15 hrs at 250 rpm.

## METHODS

### *C. elegans* immunofluorescence analysis

Immunostaining was performed as previously described with adaptations.^70^ All *C. elegans* strains were grown on NGM until the L4 to young adult stage. For Figure 6, animals were grown on NGM until the L4 stage and then transferred to “slow-kill” plates containing either 1% DMSO (solvent control) or 200 µM PYO. 30 µL of 0.2% poly-L-lysine was sandwiched between two clean positively charged slides and then allowed to air dry for 30 min. Five to ten slides were prepared for each condition. About 100-200 animals were transferred onto clean, non-seeded (no *E. coli* OP50) plates for about 15-30 min to allow for the removal of excess bacteria from the cuticle. Two flat metal blocks were then placed at the bottom of a container filled with dry ice and 100% ethanol.

On each poly-L-lysine-coated slide, 25-30 animals were picked into a 5 µL droplet of M9W. A 22 x 22 mm glass coverslip was then carefully placed on top of the animals and immediately flash-frozen on the flat metal blocks. Flash-frozen slides were stored at -80 °C until ready for immunostaining. Without thawing the slides, a razor edge was used to quickly flick off the coverslip. Slides were immersed in Coplin jars containing cold 100% methanol for 15 min at -20°C followed by 100% acetone for another 15 min at -20 °C. Slides were then air-dried at room temperature.

Animals were rehydrated with 50-100 µL 1X PBS and then blocked for 30 min with 5% normal goat serum in 1X PBS at room temperature. Primary antibodies were prepared in 5% normal goat serum in 1X PBS containing 0.2% Tween-20 (PBST) at the following concentrations: 1:250 mouse anti-FLAG M2 (for TIR-1) and 1:500 rabbit anti-PGP-2. Slides were incubated with primary antibody overnight in a humidified chamber at 4 °C. Slides were washed with PBST and then incubated with secondary antibody (anti-rabbit AF647 or anti-mouse AF555) in 5% normal goat serum in PBST for 1 hr at room temperature. Slides were washed again with PBST followed by 1X PBS. Slides were then mounted with SlowFade Gold containing DAPI (Thermo Fisher) and sealed with clear nail polish.

All immunostained slides were subsequently imaged on the THUNDER Imager (Leica DMi8, inverted microscope) using a Leica 63X objective.

### Feeding RNAi

*C. elegans* were fed *E. coli* HT115 expressing dsRNA targeting the genes of interest, as previously described with modifications.^60,71,72^ In brief, HT115 bacteria expressing specific dsRNA were grown on LB agar containing 50 µg/mL ampicillin and 15 µg/mL tetracycline at 37°C overnight. Colonies were then inoculated in LB broth containing 50 µg/mL ampicillin overnight at 37 °C for 16-18 hrs with shaking at 250 rpm. Overnight cultures were then seeded onto NGM plates containing 5 mM IPTG and 50 µg/mL carbenicillin and incubated for 16-18 hrs. Synchronized L1 animals were then transferred onto NGM plates with the grown bacteria and allowed to mature to the L4 stage.

### LysoTracker Red and LysoSensor Green assays

Animals were stained with either LysoTracker Red (Thermo Fisher) or LysoSensor Green (Thermo Fisher) as previously described.^35,40^ To stain animals with either LysoTracker Red or LysoSensor Green, 60 mm NGM plates (about 10 mL media) were first seeded with *E. coli* OP50 or HT115 and allowed to dry. Stocks were then diluted 1:10 in M9W (100 µM). 100 µL of each dye was then added on top of the dried bacteria and the dye was allowed to percolate through the plates for 1-2 hrs (final concentration 1 µM). L1 synchronized animals were then dropped on NGM plates containing either 1-2% DMSO, 1 µM LysoTracker Red, or 1 µM LysoTracker Red plus 1 µM LysoSensor Green and allowed to grow in the dark until the L4 stage. About 1-2 hrs before imaging, animals were transferred to NGM plates containing freshly seeded *E. coli* OP50. For infection or phenazine supplementation experiments, L4 animals were transferred to challenge conditions for 4-6 hrs at 25 °C before imaging.

For N-acetylcysteine (NAC) supplementation experiments (Figures 4M-4O), PGP-2::GFP animals were grown on 1 µM LysoTracker Red and then transferred to plates containing 5 mM NAC for 2 hrs at 22°C and 1 µM LysoTracker Red in the dark. Animals were then transferred onto “slow-kill” media plates containing solvent control (1% DMSO) or 200 µM PYO for 4 hrs at 25°C in the dark and imaged on the THUNDER Imager.

For Figures 4A, 4F, and 4G, animals were imaged on the THUNDER Imager (Leica DMi8, inverted microscope) using a Leica 63X objective. For Figures 4D and 4E, animals were imaged using a Zeiss AXIO Imager Z2 microscope.

### Microscopy and image analysis

#### Immunofluorescence

All immunostained slides were imaged using the THUNDER Imager (Leica DMi8, inverted microscope) with a Leica 63X objective and LasX software (Leica). Z-stacks were taken for each animal. The width of each slice was automatically determined by the software. Each stack was about 20 µm. The LasX 3D analysis software was used to quantify the volume of each PGP-2+ vesicle within each animal for the indicated conditions.

#### Live imaging of animals

For fluorescence imaging, nematodes were mounted onto 2% agarose pads, paralyzed with 50 mM tetramisole (Sigma), and imaged using a Zeiss AXIO Imager Z2 microscope with a Zeiss Axiocam 506 mono camera and Zen 2.5 (Zeiss) software. For LysoTracker Red and/or LysoSensor Green, or TIR-1::wrmScarlet imaging, animals were mounted onto 2% agarose pads, paralyzed with 50 mM tetramisole (Sigma), and sealed with coverslips and VALAP. Slides were then inverted and imaged on the THUNDER Imager (Leica DMi8, inverted microscope) with a Leica 63X objective and LasX software (Leica).

#### TIR-1/SARM1 puncta quantification

TIR-1/SARM1 puncta quantification was performed as described previously. ^12^

#### Co-localization of TIR-1/SARM1 and PGP-2 in live animal samples

Co-localization of TIR-1/SARM1 and PGP-2 in live animals was determined using Fiji (ImageJ). First, red (TIR-1) and blue (autofluorescence) channels were co-localized with the “Color threshold” function. Green (PGP-2 or LMP-1) and blue (autofluorescence) channels were then co-localized. Any co-localized signal with the blue channel was then subtracted from the images. JACoP was then used to determine the Pearson’s coefficient of co-localization between red and green channels.^73^

#### Co-localization of TIR-1/SARM1 with PGP-2+ vesicles in immunostained samples

Aivia (Leica) was used to quantify co-localization between TIR-1/SARM1 and PGP-2+ vesicles in the immunostained samples. Briefly, pixel classifiers were used to determine positive signals for both TIR-1/SARM1 (Alexa Fluor 555) and PGP-2 (Alexa Fluor 647) channels. Meshes were then made for each channel and the number of relations were determined between TIR-1/SARM1 and PGP-2. The 3D object analysis recipe for TIR-1/SARM1 was as follows: Image Smoothing Filter Size: 1; Min Edge Intensity: 50; Fill Holes Size: 0; Object radius: 0-2 µm; Mesh smoothing factor: 0; Min Edge to Center Distance: 0.5 µm. The 3D object analysis recipe for PGP-2 was as follows: Image Smoothing Filter Size: 1; Min Edge Intensity: 70; Fill Holes Size: 0; Object radius: 0-5 µm; Mesh smoothing factor: 0; Min Edge to Center Distance: 0.5 µm. Relations on frame were then quantified and graphed with the PGP-2 channel as the primary mesh.

#### TIR-1/SARM1 puncta size on PGP-2+ lysosome-related organelles

For quantification of TIR-1/SARM1 puncta size on lysosome-related organelles, Aivia (Leica) was used to first render the Z-stacks imaged from co-immunostained animals treated with solvent control or pyocyanin in three dimensions. For lysosome-related organelles that co-localized with TIR-1/SARM1, we quantified the size (volume) of all TIR-1/SARM1 objects associated with that vesicle.

Quantifications were completed in 5 animals, where 5 random PGP-2+ vesicles in the anterior intestine per animal were chosen.

#### *C. elegans* strain construction

All CRISPR genome editing was performed as previously described.^74^ CRISPR-Cas9 editing with single-stranded oligodeoxynucleotide (ssODN) homolog-directed repair was used to introduce the hydrophobic domain deletion into *tir-1.* All CRISPR reagents were purchased from Integrated DNA Technologies. Target guide sequences were selected using the CHOPCHOP web tool. The ssODN repair templates contained indicated deletions with 35 bp flanking homology arms. ssODN sequences are listed in Table S4. The F1 progeny were screened for Rol phenotypes 3 to 4 days after injection and then for indicated edits using PCR and Sanger sequencing. Primer sequences used for genotyping are listed in Table S4.

### Immunoblot analyses

Protein lysates were prepared using a Teflon Dounce homogenizer from 5-10,000 *C. elegans* grown to the L4 larval stage on NGM plates seeded with *E. coli* OP50, as previously described ^12,20^. LDS Sample Buffer (Thermo Fisher Scientific) was added to a concentration of 1X with 5% β-mercaptoethanol. All samples were incubated at 70 °C for 10 minutes. Total protein from each sample was resolved on NuPage Bis-Tris 4–12% gels (Invitrogen), for detection of phosphorylated and total p38 PMK-1, or NuPage Tris-Acetate 3-8% gels (Invitrogen), for detection of TIR-1. Protein was then transferred to 0.2 µM nitrocellulose membranes (Bio-Rad) and blocked with 5% milk in 1x TBS + 0.2% Tween-20 for one hour. Blots were then probed with a 1:1000 dilution of mouse monoclonal anti-FLAG M2 (Sigma, #F1804), 1:1000 phospho p38 PMK-1 (Cell Signaling Technology, #9211), 1:1000 total PMK-1,^45^ 1:2000 mouse monoclonal anti-alpha-Tubulin (Sigma, #T5168), overnight at 4 °C. Anti-mouse IgG-HRP (Abcam, #ab6789) or anti-rabbit IgG-HRP (Cell Signaling Technology, #7074) secondary antibodies were used at a dilution of 1:10,000 to detect the primary antibodies. Blots were then developed with the addition of SuperSignal™ West Pico or West Femto PLUS Chemiluminescent Substrate (Thermo Fisher Scientific) and visualized using a ChemiDoc MP Imaging System (Bio-Rad). Band intensities were quantified using ImageJ (Fiji).

NativePAGE analysis was performed as previously described. Protein lysates were prepared in 1x NativePAGE buffer (Invitrogen), 1% digitonin, and HALT protease inhibitor (Thermo Fisher). Coomassie G-250 additive was added to samples and loaded onto NativePAGE 3-12% Bis-Tris gels (Invitrogen). Samples were first run in dark blue cathode buffer (inner chamber) until the front ran 1/3 down the gel. The buffer was then switched to the light blue cathode buffer. Protein was then transferred onto 0.2 µm PVDF membranes overnight at 4°C. After transfer, membranes were placed in 8% acetic acid and fixed for 15 minutes at room temperature on a rocker. The blot was then destained with 50% methanol/10% acetic acid until the membrane was white and blocked with 5% milk in 1x TBS + 0.2% Tween-20 for 1 hr at room temperature on a rocker. Blots were then transferred to mouse anti-FLAG M2 (Sigma, #F1804) antibody overnight at 4°C. The following day, blots were washed and transferred to anti-mouse IgG-HRP (Abcam, #ab6789) for 1 hr at room temperature on a rocker. Blots were then developed with the addition of SuperSignal^TM^ West Femto PLUS Chemiluminescent Substrate (Thermo Fisher Scientific) and visualized using a ChemiDoc MP Imaging System (Bio-Rad).

### qRT-PCR analyses

RNA was extracted from about 2000 L4 animals and reverse transcribed to cDNA using the iScript cDNA Synthesis Kit (Bio-Rad). cDNA was then analyzed on the CFX384 thermocycler (Bio-Rad) with primers described in Table S4. All values were normalized to the geometric mean of the control genes *act-3* and *snb-1*. Relative expression was then calculated using the Pfaffl method.^75^

### *C. elegans* pathogenesis assays

“Slow-killing” *P. aeruginosa* infection experiments were performed as previously described.^76^ In brief, *P. aeruginosa* was grown as described above and 10 µL overnight culture was spread onto the center of 35-mm tissue culture plates containing 4 mL slow-kill agar (0.35% Bacto-peptone, 0.3% sodium chloride, 1.7% agar, 5 µg/mL cholesterol, 25 mM potassium phosphate, 1 mM magnesium sulfate, 1 mM calcium chloride). Plates were then incubated for 24 hrs at 37°C followed by 24 hrs at 25 °C. Wild-type and *pgp-2(kx48) C. elegans* were then transferred to *P. aeruginosa* slow-kill plates containing FUDR. Dead animals were scored twice daily until completion. Three trials of the assay were performed. Sample sizes, mean survival, and p-values for all trials are shown in Table S1.

“Fast-killing” *P. aeruginosa* infection experiments were performed as previously described.^20,46,76^ Briefly, PA14 Δ*phz* mutants were grown on PGS agar (1% Bacto-peptone, 1% glucose, 1% NaCl, 150 mM sorbitol, 1.7% Bacto-agar) for 24 hrs at 37 °C followed by 24 hrs at 25 °C. The lawns were then scraped off of the agar. The remaining agar was then melted, and the pH was adjusted to pH 7.0 using 100 mM potassium phosphate. Either DMSO (solvent) or 50 µM PYO was then added to the agar. Once cooled and solidified, plates were seeded with 10X *E. coli* OP50. Wild-type, TIR-1^ΔHD^ mutants or *pgp-2(kx48)* mutants were then transferred to the plates at the L4 stage and scored for survival at room temperature (22 °C).

### Quantification, statistical analysis, and visualization

Differences in the survival of *C. elegans* in the *P. aeruginosa* pathogenesis assays were determined with the log-rank test after survival curves were estimated for each group with the Kaplan-Meier method. OASIS 2 was used for these statistical analyses.^68^ Statistical hypothesis testing was performed with Prism 9 (GraphPad Software) using methods indicated in the figure legends. Table S2 contains all source data and statistical analysis methods and results. Sample sizes, survival, and p-values for all trials are shown in Table S1. BioRender was used to create the image in Figure 6.

## SUPPORTING INFORMATION

**Figure S1.**
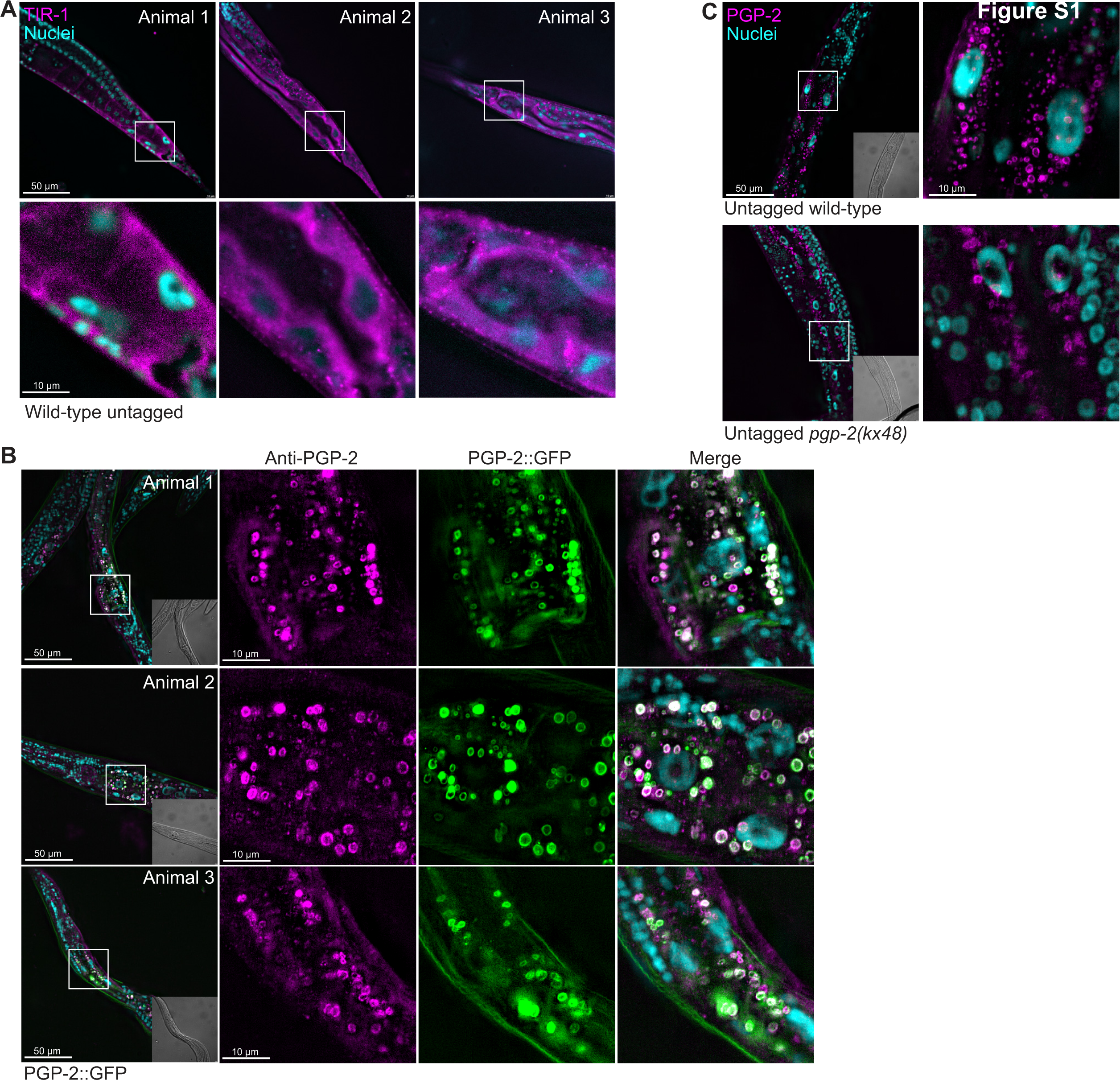
TIR-1/SARM1 is expressed on lysosome-related organelles in *C. elegans* intestinal epithelial cells. **(A)** Three representative images of untagged animals immunostained with anti-FLAG (for TIR-1) antibody. Dotted white boxes indicate higher magnifications. **(B)** Three representative images of PGP-2::GFP animals immunostained with anti-PGP-2 antibody. Insets in left panel represent corresponding DIC images. **(C)** Representative images of wild-type and *pgp-2(kx48)* loss-of-function mutants immunostained with anti-PGP-2 antibody. Dotted white boxes indicate higher magnifications. Insets in left panel represent corresponding DIC images. Scale bars as indicated. Related to Fig. 1.

**Figure S2.**
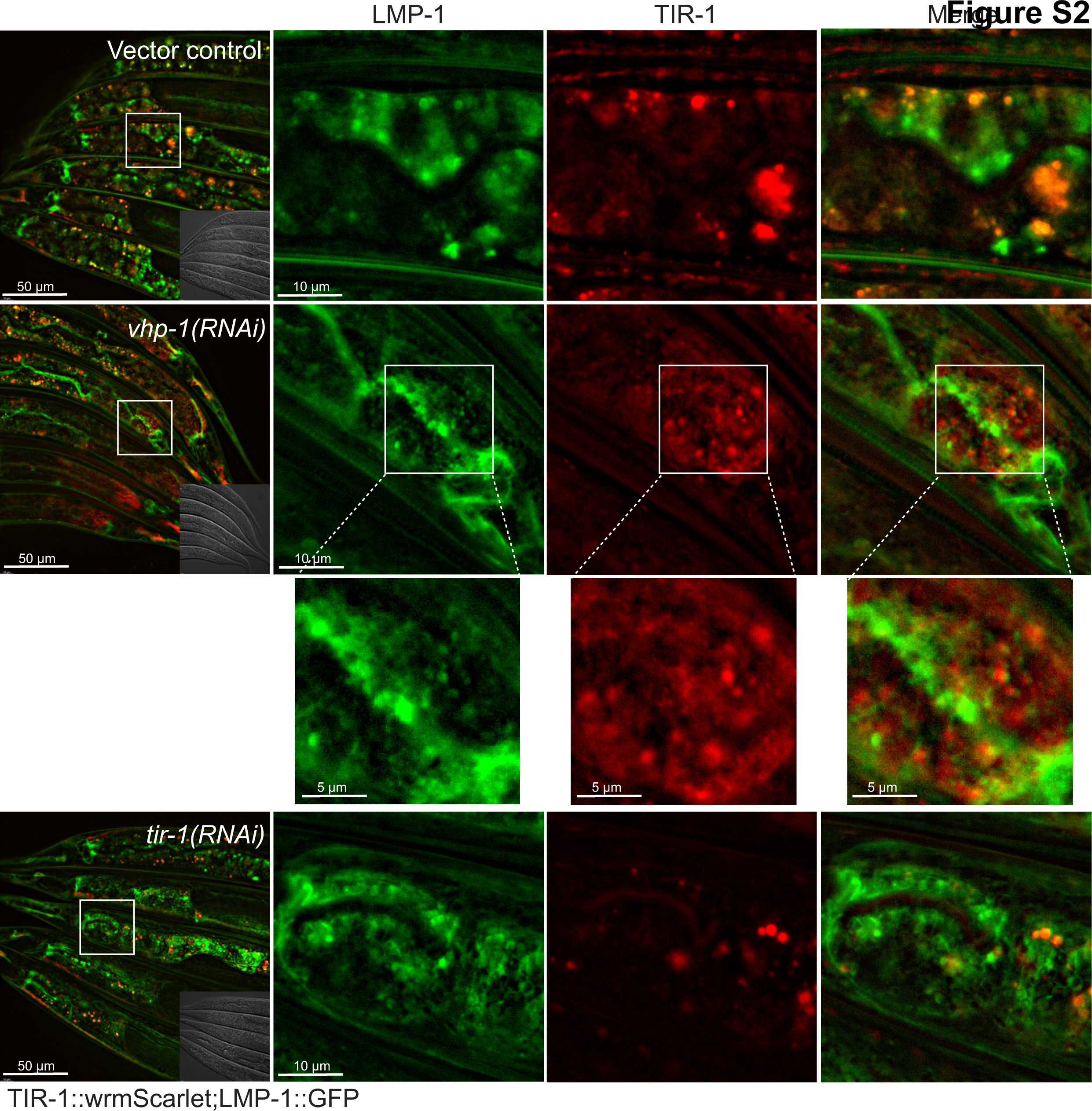
TIR-1/SARM1 co-localization with lysosome-related organelles can be visualized in live *C. elegans*. Images of *C. elegans* TIR-1::wrmScarlet animals crossed into a LMP-1::GFP translational reporter and treated with either vector control, *vhp-1(RNAi)* or *tir-1(RNAi)*. Dotted white boxes indicate higher magnifications. Scale bars as indicated. Related to Fig. 2.

**Figure S3.**
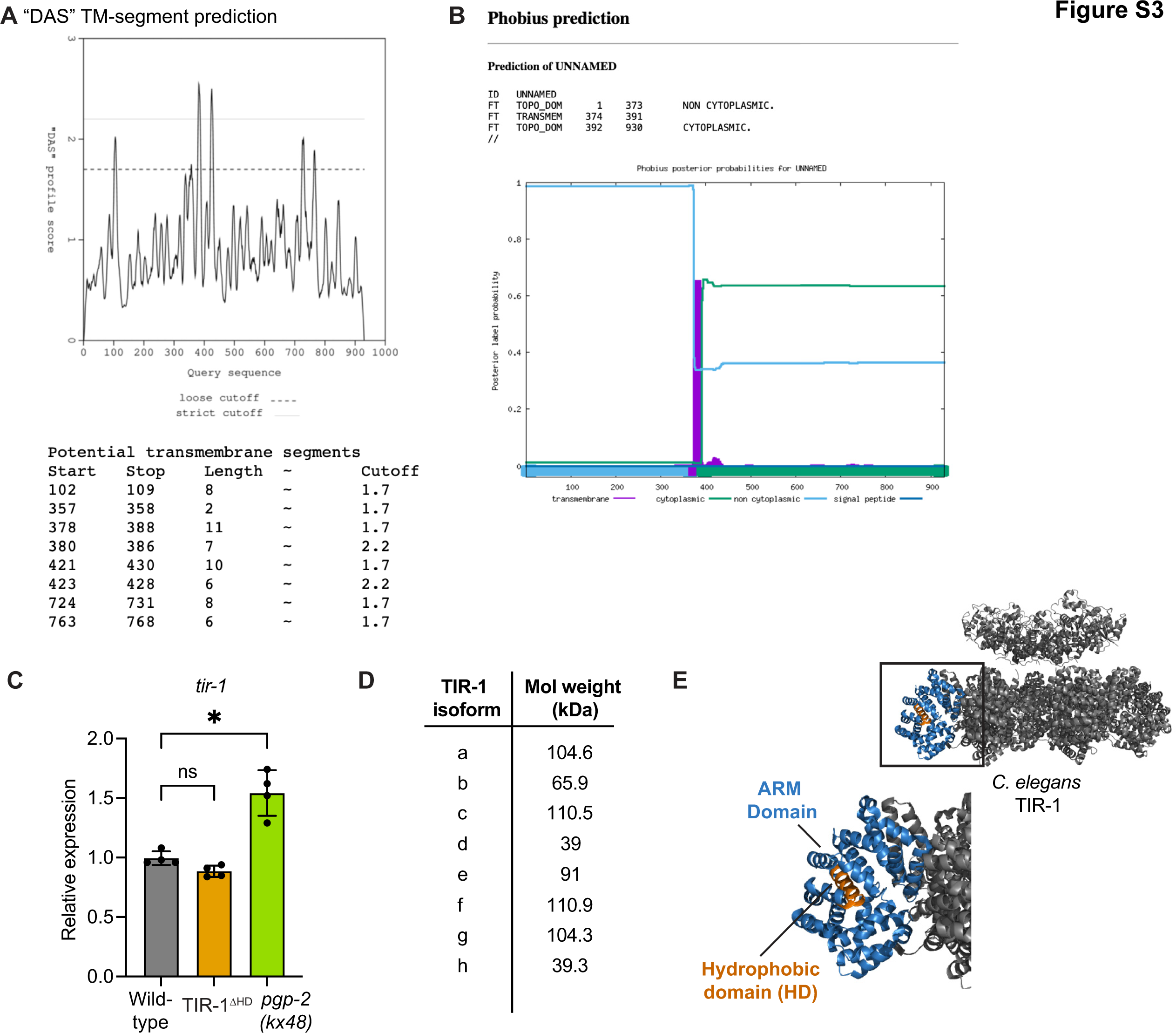
A hydrophobic domain in the TIR-1/SARM1 ARM domain is required for its localization on the membranes of lysosome-related organelles and p38 PMK-1 activation. **(A)** Prediction of hydrophobic domains in TIR-1a by DAS. **(B)** Prediction of hydrophobic domains by Phobius. **(C)** qRT-PCR of *tir-1* in wild-type, TIR-1^ΔHD^, or *pgp-2(kx48)* mutants. *equals p<0.05 (one-way ANOVA with Dunnett’s multiple comparisons test). **(D)** Predicted molecular weights of TIR-1/SARM1 isoforms from WormBase. **(E)** Predicted structure of the hydrophobic domain in *C. elegans* TIR-1/SARM1.^39^ Related to Fig. 3.

**Figure S4.**
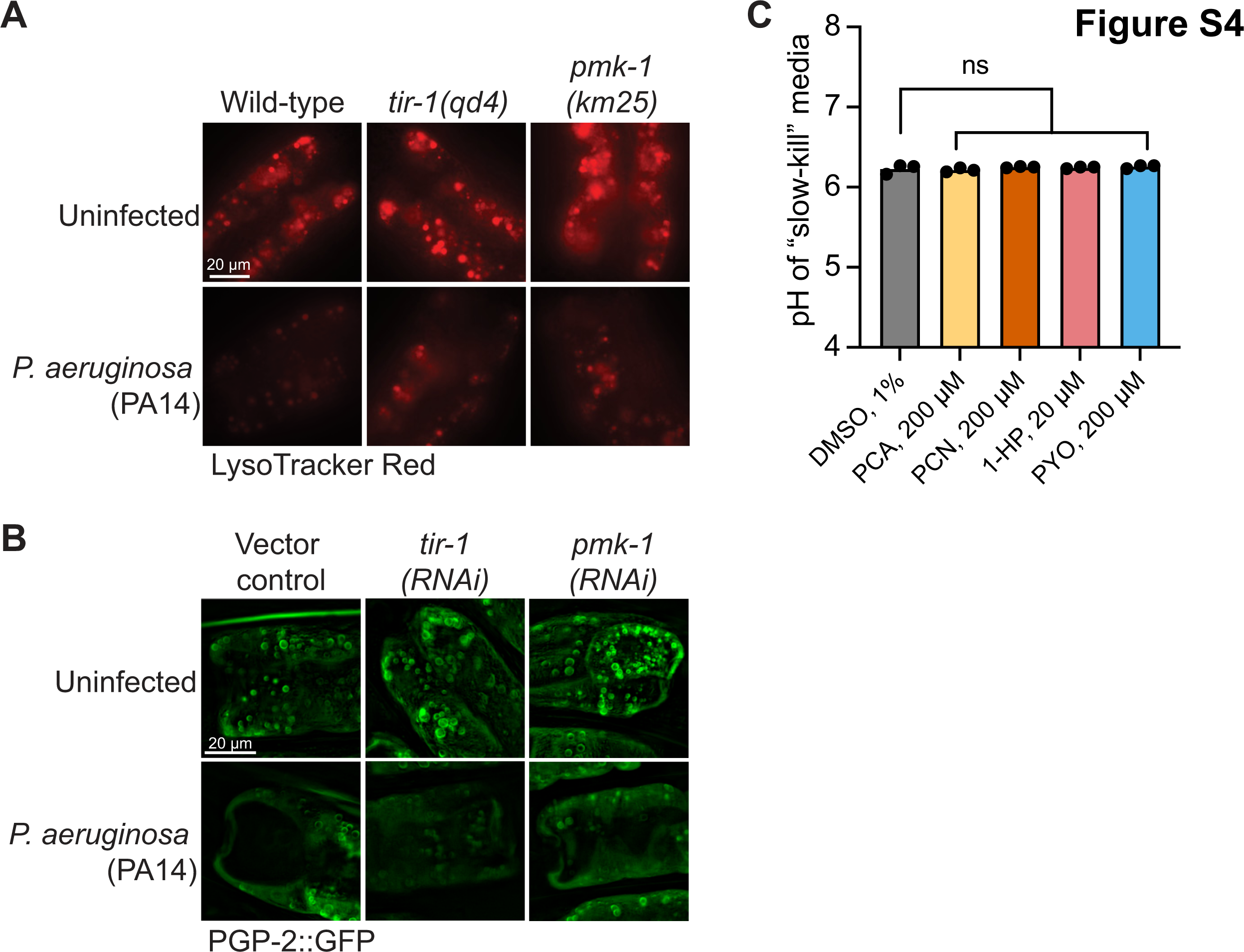
The pathogen-derived virulence effector pyocyanin alkalinizes and condenses lysosome-related organelles in intestinal epithelial cells. **(A)** Representative images of wild-type, *tir-1(qd4)* and *pmk-1(km25)* animals stained with LysoTracker Red and uninfected or infected with *P. aeruginosa* (PA14). **(B)** Representative images of PGP-2::GFP animals treated with vector control, *tir-1(RNAi)* and *pmk-1(RNAi)* infected with *P. aeruginosa.* (C) pH of liquid “slow kill” media supplemented with indicated phenazines. Related to Fig. 4.

**Figure S5.**
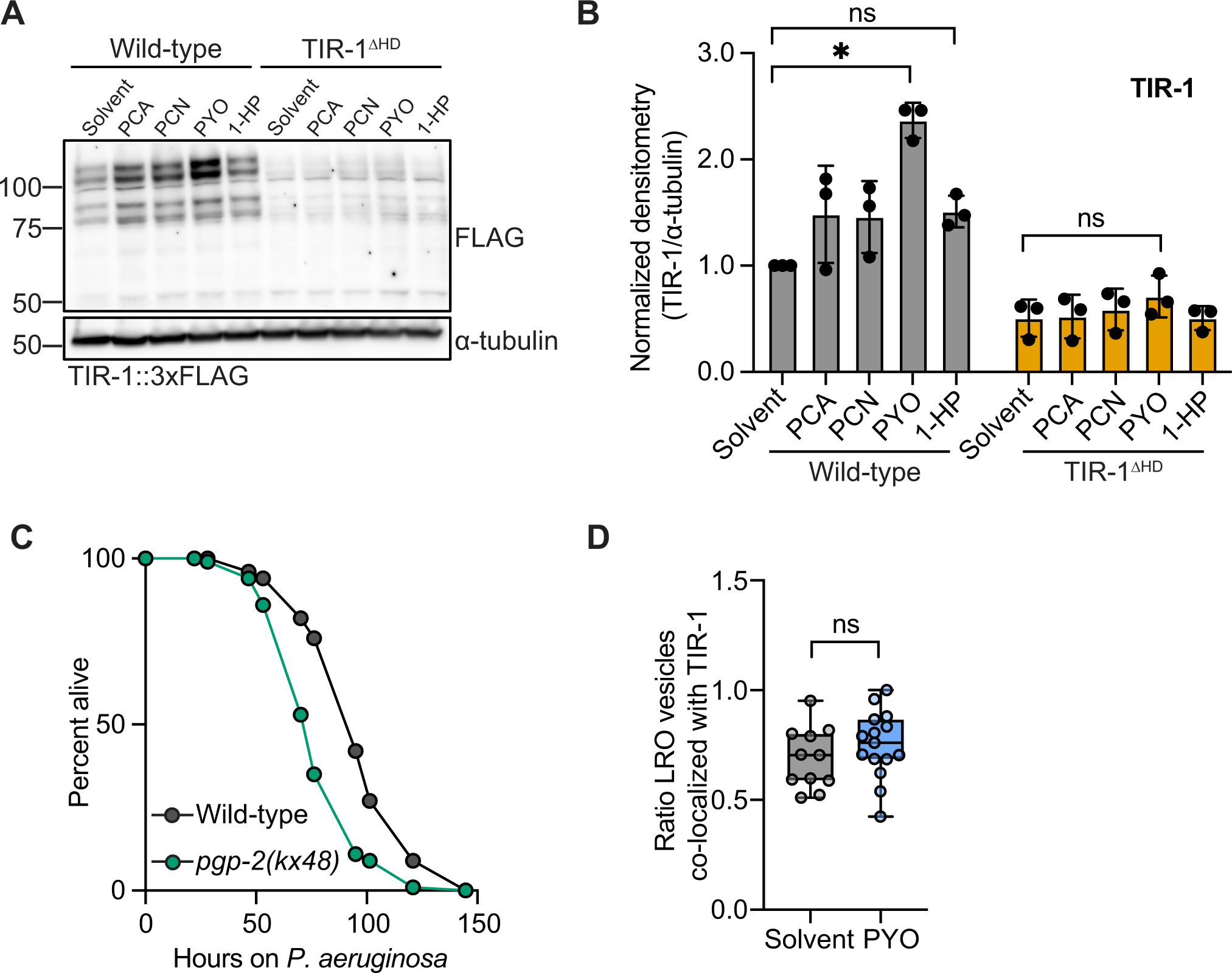
Pyocyanin triggers aggregation of TIR-1/SARM1 on lysosome-related organelles and activates p38 PMK-1 intestinal immunity. **(A)** Representative immunoblot of whole cell lysates isolated from wild-type and *pgp-2(kx48)* mutants treated with solvent control or PYO for 2-6 hrs probed with anti-phosphorylated PMK-1, anti-total PMK-1, and anti-⍺-tubulin. **(B)** Densitometric quantification of conditions in (A) (*n=3).* *equals p<0.05 (two-way ANOVA with Šídák’s multiple comparisons test). **(C)** Representative *P. aeruginosa* pathogenesis assay on wild-type vs. *pgp-2(kx48)* mutants. The difference between wild-type and *pgp-2(kx48)* is significant (p<0.05, log-rank test) (n=3). **(D)** Quantification of the ratio of PGP-2+ vesicles co-localized with TIR-1/SARM1 from co-immunostained samples in the presence or absence of PYO. ns=not significant. Mean lifespans and statistics for all replicates are in Table S1. Related to Fig. 5.

**Table S1.** Sample sizes, mean lifespans, and p-values for the *C. elegans* pathogenesis assays.

**Table S2.** Source data and statistical tests used for each figure and supplemental figure.

**Table S3.** *P. aeruginosa* transposon mutant hits that rescued LysoTracker Red phenotype.

**Table S4.** Primer, crRNA guide and ssODN sequences designed for this study.

**Video S1.** Three-dimensional reconstruction of co-immunostained anti-FLAG (TIR-1) and anti-PGP-2 animal.

